# MRSA emerges through natural transformation

**DOI:** 10.1101/2020.12.08.415893

**Authors:** Mais Maree, Le Thuy Thi Nguyen, Ryosuke L. Ohniwa, Shenghe Huang, Masato Higashide, Tarek Msadek, Kazuya Morikawa

## Abstract

Methicillin-resistant *Staphylococcus aureus* (MRSA) carries the resistance gene *mecA* in the staphylococcal cassette chromosome (SCC) that disseminates among staphylococci but the cell-to-cell transmission mechanism of SCC has not been clarified for half a century^1^. Here, we present evidence for efficient natural transformation in *Staphylococcus aureus* and its relevance in SCC*mec* transmission. We found that growth in biofilm conditions increased the transformation efficiency in a dependent manner on two component signal transduction systems, TCS13 (AgrCA) and TCS17 (BraSR). Strikingly, we demonstrate that natural transformation mediates the transfer of SCC*mec* from MRSA or methicillin-resistant coagulase negative staphylococci to methicillin-sensitive *S. aureus*. The site-specific insertion/excision system mediated by cassette chromosome recombinases was essential for SCC*mec* transformation while the stability of SCC*mec* varied depending on SCC types and recipients. We propose that natural transformation is the key process in the emergence of MRSA.

## Introduction

*Staphylococcus aureus* is a Gram-positive bacterium belonging to Phylum *Firmicutes* that is resident in the nasal cavities of about 30 percent of the entire population. These *S. aureus* carriers are normally asymptomatic but opportunistic infections, ranging from minor skin abscesses to severe diseases (such as pneumonia, osteomyelitis, or toxic shock syndrome), are possible. Immunocompromised hosts are vulnerable but the spread of community-associated infections by highly virulent *S. aureus* has also been reported^2^.

Antibiotic resistance is the most notorious feature of this pathogen, especially methicillin-resistant *S. aureus* (MRSA). MRSA is the leading cause of nosocomial infections (health-care-associated MRSA; HA-MRSA) and is also associated with healthy individuals (community-associated MRSA; CA-MRSA) and livestock (livestock associated MRSA; LA-MRSA), posing a global health burden^3,4,5,6^. The percentage of MRSA among *S. aureus* isolates is diverse among countries (Vietnam 73%, United States 45%, Japan 41%, North Europe 1%), raising concerns in clinics, care homes and other places with high densities of immunocompromised individuals^7^.

The global spread of this major human pathogen can be explained by an arsenal of virulence factors and antibiotic resistance genes, many of them located on mobile genetic elements (MGEs) such as plasmids, prophages, transposons, pathogenicity islands, insertion sequences, and the staphylococcal cassette chromosome (SCC)^8,9^. In MRSA and methicillin-resistant coagulase negative staphylococci (MR-CoNS), the methicillin-resistant determinant *mecA* is always located within the SCC (SCC*mec*). SCC*mec* is itself a 20-60 kb genetic element integrated by Ccr recombinases at a specific site (*attB*) in *orfX* near the replication origin of the chromosome^10^. Epidemiological studies show that these SCC*mec* elements are transmitted among staphylococci and at least 20 independent acquisitions of SCC*mec* were reported to have occurred in *S. aureus*^11^, but the exact mechanism of cell-to-cell transmission has been debated for over 50 years^1^ (see Discussion).

The presence of diverse MGEs conveying virulence and resistance factors to the *S. aureus* genome indicates a prominent evolutionary ability mediated by horizontal gene transfer (HGT). Bacteriophage-mediated transduction and conjugative machinery-dependent conjugation are historically well-characterized HGT mechanisms in staphylococci, with the former considered to be the primary method^1^. In 2012, a subpopulation of *S. aureus* was shown to develop natural genetic competence for DNA transformation by expressing competence machinery (DNA incorporation machinery) genes in the *comG* and *comE* operons, which are under the direct transcriptional control of the ‘cryptic’ sigma factor SigH^12,13^. Bacteria modified to overexpress SigH incorporated plasmid DNA as well as SCC*mec* II elements ^13^. However, the transformation efficiencies of the unmodified model strains (N315 or N315ex) and other clinical isolates in that report were under the detection limit (<10^−11^)^13^. Competence transcription factor ComK was also shown to synergistically upregulate many competence genes^14^. However, efforts to transform the tested strains by overexpressing SigH and ComK was unsuccessful^14^. These observations have led to the current belief that natural transformation plays no role in staphylococcal evolution, including the multiple, independent emergence of MRSA strains with diverse genetic backgrounds^15^.

In the present study, we identify specific two-component systems (TCSs)^16,17^ involved in the regulation of the competence operon promoter (P_*comG*_). As TCSs are major mediators of sensing and environmental response, we conducted a survey to clarify the conditions that trigger efficient transformation in *S. aureus* through this mechanism. Strikingly, we present the first experimental evidence of efficient inter- and intraspecies transfer of SCC*mec* among staphylococci. We propose natural transformation as a major evolutionary strategy for this ever-adaptive pathogen.

## Results

### TCSs are involved in the expression of *comG* promoter in subpopulations

To delineate conditions conducive to natural transformation, we generated a series of

15 TCS deletion mutants, removing each TCS (Δ3~Δ17) except the essential TCS1 (WalKR)^18^, in the *S. aureus* strain N315ex w/o ϕ^13^ (termed Nef; Supplementary Table 1) (Fig. 1a). Nef is an N315 derivative strain that can develop natural genetic competence but does not possess any conjugative elements or a lysogenic phage that transfers DNA by transduction or pseudo-competence. Nef also lacks the SCC*mec* and its embedded TCS2 (*SA0066-SA0067*).

**Figure 1.**
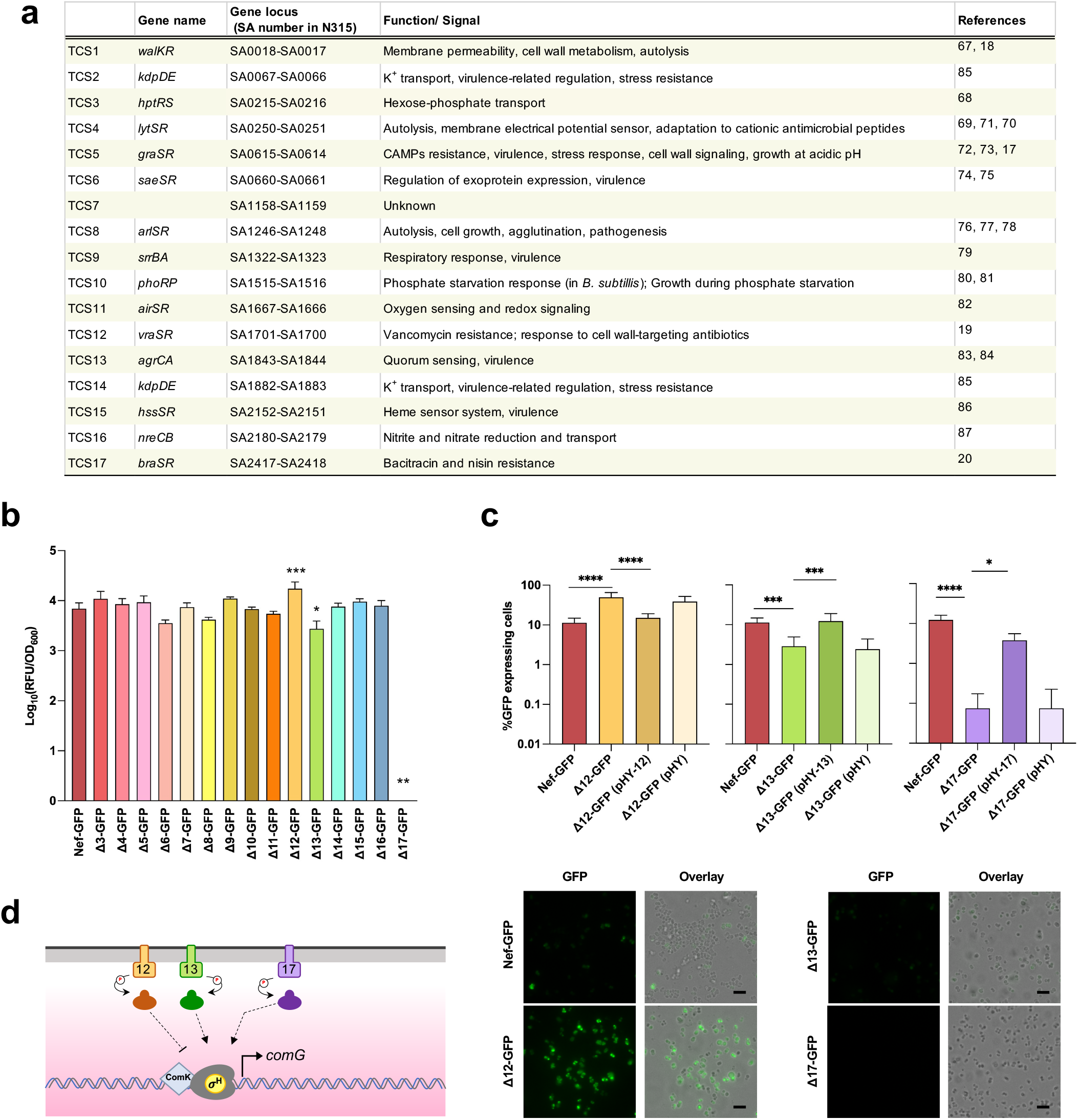
TCS12, TCS13 and TCS17 are involved in *comG* promoter activity. **a**, 17 TCSs in *S. aureus* N315. Gene name and locus in N315 genome are shown. b,c, ΔTCS mutants (Δ3-Δ17) were tested for involvement in the regulation of P_*comG*_. Nef and its derivative ΔTCS carrying the P_*comG*_-*gfp* reporter were grown in CS2 medium with shaking. **b**, The Y axis shows the increase in RFU/OD_600_ values, which was calculated by subtracting the minimum GFP intensities from the maximum GFP intensities during 12-24 h of growth. The mean of n=3-8 independent experiments are shown. Error bars represent s.d. Statistical significance was determined by one-way ANOVA with Dunnett’s multiple comparison test. * P < 0.05, ** P < 0.01, *** P < 0.001. **c**, The population percentage expressing GFP was determined after 12-14 h of growth by fluorescent microscopy (**bottom**). The mean of n=4-9 independent experiments are shown. Error bars represent s.d. Statistical significance was determined by one-way ANOVA with Tukey’s multiple comparison test. * P < 0.05, ** P < 0.01, *** P < 0.001, ^****^ P < 0.0001. Scale bars, 5 μm. d, Schematic summary of the TCSs involved in P_*comG*_ regulation.

The activity of the SigH-dependent *comG* operon promoter (P_*comG*_) was monitored by a GFP reporter (P_*comG*_-*gfp* transcriptional fusion) in every ΔTCS. In planktonic cultures using CS2 medium, the GFP intensity of the parental strain (Nef-GFP) initially increases from 8 h and peaks around 15 h (Supplementary Fig. 1a). In contrast, no GFP could be detected in other media such as TSB (Supplementary Fig. 1b), in line with our previous observations that the activation of the *comG* promoter is dependent upon culture conditions^13^.

The ΔTCSs did not exhibit major growth defects in TSB (Supplementary Fig. 2a). However, in CS2 medium, Δ5, Δ12, and Δ13 exhibited minor growth defects compared with Nef while Δ9 and Δ17 exhibited a higher yield (Supplementary Fig. 2b). Figure 1b shows the peak values of the reporter expression (intensities of GFP fluorescence per OD) in ΔTCSs cultured in CS2 medium. Compared with the parental strain, both Δ13 and Δ17 showed significantly lower values while Δ12 achieved a higher value and other ΔTCSs had no significant effect. We also used fluorescence microscopy to observe cell populations expressing the GFP reporter (Fig. 1c) and, in the parental strain, 11.3% of the cells expressed GFP after culturing for 12-14 h in CS2 medium but, in Δ13 and Δ17, only 2.9% and 0.1% of the cells expressed GFP (Fig. 1c, middle and right panels). In Δ12, on the other hand, 49.3% of the cells expressed GFP, an approximately 4-fold higher expression than the parental strain (Fig. 1c, left panel). Complementation *in trans* within these ΔTCSs restored the percentages of GFP-positive cells towards the parental strain levels. Taken together, our results suggest the involvement of TCS12, TCS13, and TCS17 in the regulation of natural genetic competence in Nef (Fig. 1d).

**Figure 2.**
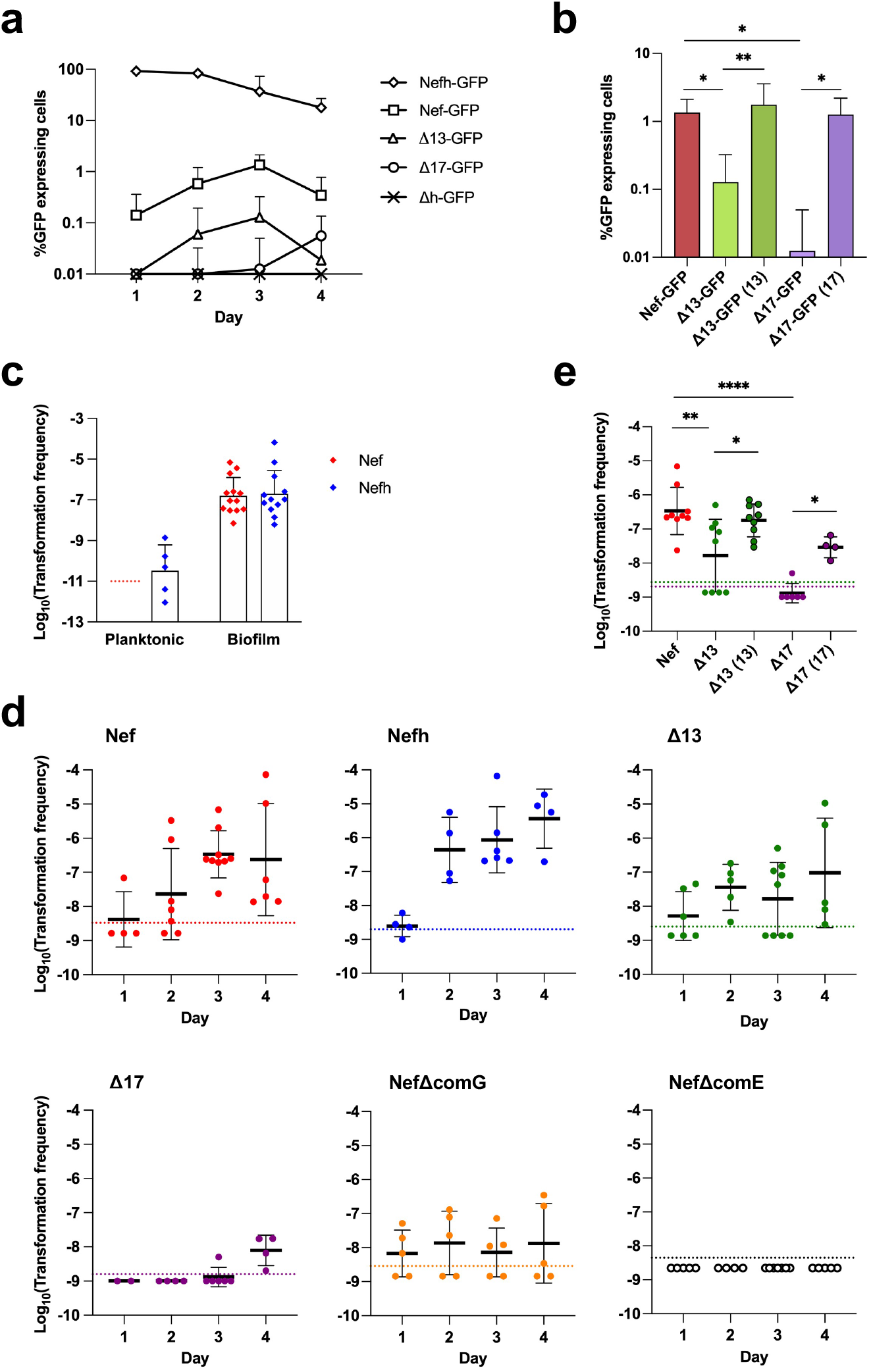
TCSs 13 and 17 are important for *comG* promoter activity and natural transformation in biofilm growth conditions. **a,** The percentage of Nef and its derivatives expressing P_*comG*_-*gfp* reporter. Cells were statically grown in CS2 medium. The mean n=3-10. Error bars represent s.d. **b,** Chromosomal complementation of the mutants restores the percentage of cells expressing GFP at day 3 in the biofilm. The mean n=7-10. Error bars represent s.d. **c,** Transformation frequencies of Nef (red) and Nefh (blue) at day 3 in the biofilm growth condition compared with the planktonic growth condition. Transformants were selected by tetracycline. Dotted line represent detection limit of planktonic Nef. **d,** Time-course development of natural transformation in the biofilm. Nef and its derivatives were statically grown in CS2 medium. Natural transformation frequencies were determined every 24 h. Dotted lines represent the detection limit. Data points represent independent experiments. Error bars represent s.d. **e,** Complementation of the TCS13 and TCS17 restores the transformation frequencies at day 3 in the biofilm. In this experiment, chromosomally complemented strains were used because plasmid-based complimented strains are tetracycline resistant. Data points represent independent experiments. Error bars represent s.d. Statistical significance (**b, e**) was determined by one-way ANOVA with Tukey’s multiple comparison test. *P<0.05, **P<0.01, ****P<0.0001.

### Cell wall-targeting antibiotics and biofilm growth conditions modulate *comG* expression

TCS12 (VraSR) is mainly involved in the response and resistance to vancomycin, but also to bacitracin and other antibiotics to some extent^19,20,21,22^, while TCS17 (BraSR) is involved in resistance to bacitracin and nisin^20^. Indeed, we confirmed that Δ12 is susceptible to vancomycin and Δ17 is susceptible to bacitracin and nisin in CS2 and TSB media (Supplementary Table 3).

In addition to their essential roles in resistance to cell wall-targeting antibiotics, TCS17 has pleiotropic roles in cell physiology, including biofilm formation^23^, and TCS13 (AgrCA) is a part of the accessory gene regulator (Agr) quorum sensing system that regulates the expression of multiple virulence genes^24^ by the diffusion sensing mechanism^25,26^ and is also involved in biofilm regulation. We noted that Δ13 and Δ17 were impaired in rigid biofilm formation compared to Nef when cultured in CS2 medium (Supplementary Fig. 3) but this phenotype is not due to growth defects as the colony forming units of Δ13 and Δ17 were not significantly different compared to Nef after 24 h under these static conditions (Supplementary Fig. 3b, right). Based on these results, we tested the effects of cell wall-targeting antibiotics and biofilm-forming, static growth conditions on P_*comG-gfp*_ expression.

**Figure 3.**
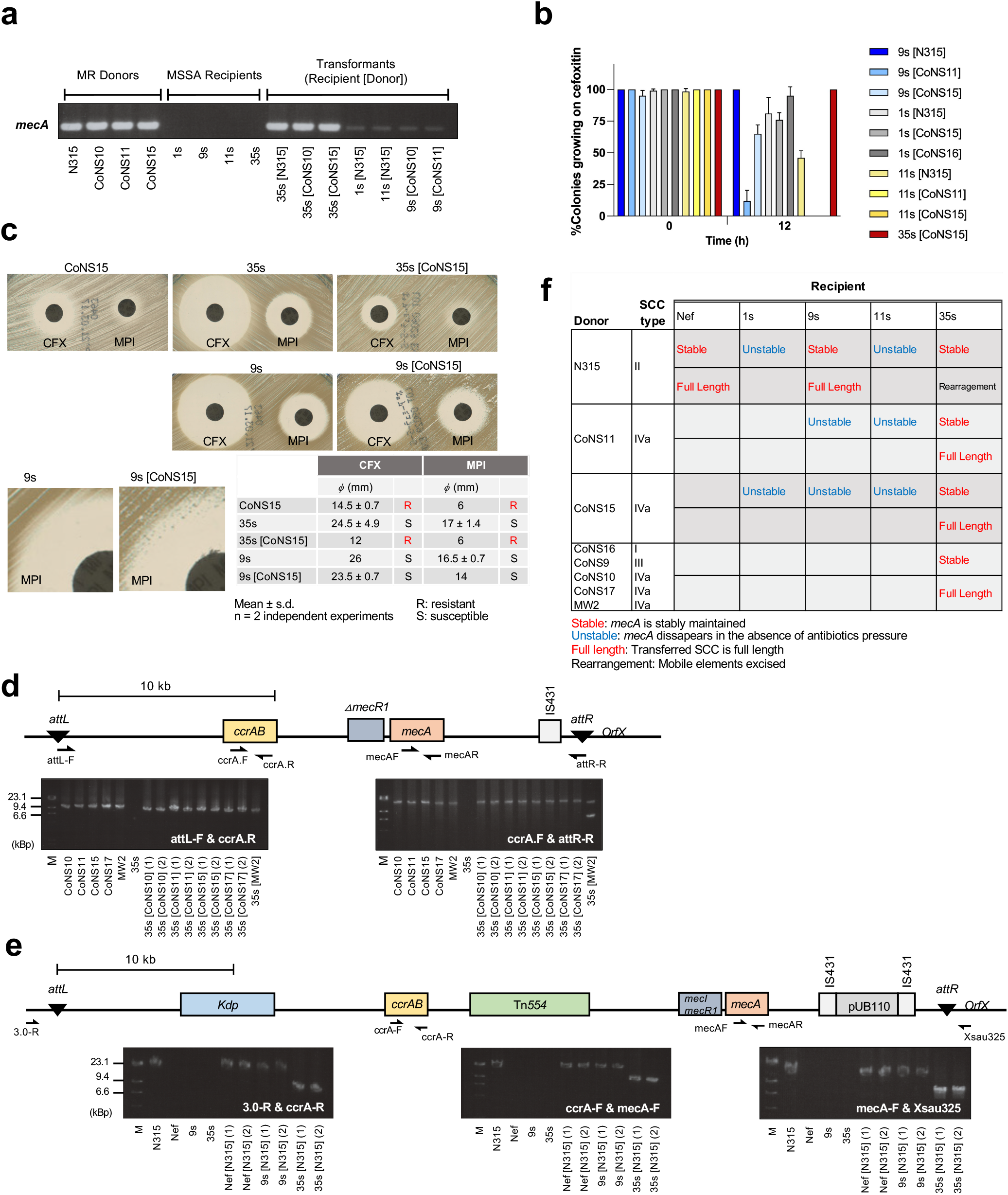
SCC*mec* transformation. **a,** Presence of the *mecA* gene was verified by PCR using mecAF and mecAR primers. Chromosomal DNA of donor and recipients were used for positive and negative controls. **b,**Stability of resistance. The transformants were passaged in drug-free media for 12 h after growth with cefoxitin (4 μg mL^−1^) for 12h (Time 0). The percentage of cells that can grow on cefoxitin was calculated by a replica method. The mean of n = 2 independent experiments is shown. Error bars represent s.d. **c,** Disk diffusion test of β-lactam antibiotics. CFX: cefoxitin, MPI: oxacillin. **right bottom**: diameters of inhibitory zones. **d,e,** Schematic structures of SCC*mec* IVa in MW2 chromosomal DNA (**d**) and of SCCme II in N315 chromosomal DNA (**e**). Primer locations are indicated with arrows. Chromosomal DNA of MR-donors and MS-recipients were used for positive and negative controls. Suffix (1), (2) represents transformants obtained from two independent experiments. (**d-e, bottom**) Long PCR verification of the entire SCC*mec* IVa (**d**) or II (**e**) elements in all transformants where *mecA* was present. M: λ *Hin*dIII. **f**, Intactness and stability of SCC*mec* in transformants.

Treatment of Nef-GFP with subinhibitory concentrations of vancomycin or bacitracin reduced P_*comG*_-*gfp* reporter expression in a concentration-dependent manner (Supplementary Figs. 4a,b). On the other hand, 8 or 16 μg mL^−1^ nisin slightly, but reproducibly, increased reporter expression intensity in Nef-GFP (Supplementary Fig. 4c).

**Figure 4.**
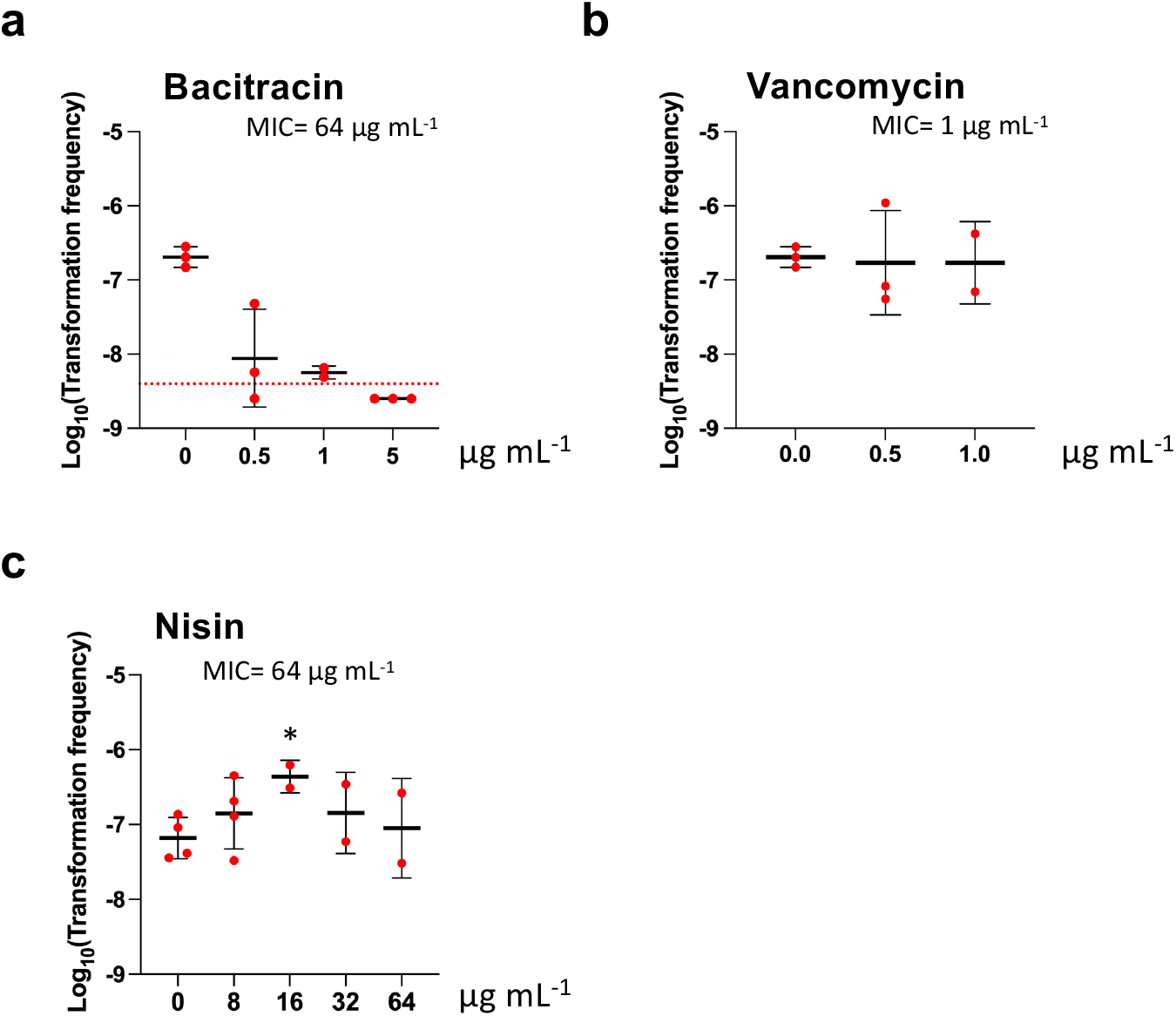
Bacitracin blocks natural transformation. Cells were grown with bacitracin (**a**), vancomycin (**b**), or nisin (**c**) under biofilm conditions in CS2 medium. Transformants were counted after 3 days. Error bars represent s.d. Dotted line represents the detection limit. Statistical significance was determined by Student’s t-test. *P<0.05.

To test bacteria under biofilm growth conditions, we cultured Nef and its derivative strains statically in CS2 medium to let cells sediment at the bottoms of a 6-well polystyrene plate where they stably attach by forming a biofilm (Supplementary Fig. 3b). In the Nef-GFP reporter strain, the percentage of GFP-expressing cells increased towards days 2-3 (Fig. 2a), an effect that was impaired in Δ13 and Δ17. The percentage of GFP-expressing cells in biofilm was less than in planktonic culture, but our data indicate that the Agr quorum sensing TCS13 plays a role in *comG* reporter expression under biofilm-forming conditions. Furthermore, we examined if other TCSs are involved in P_*comG*_-*gfp* expression under these same conditions and found that only Δ13 and Δ17 had significantly reduced GFP expression compared with Nef after 3 days (Supplementary Fig. 5) whereas Δ7, Δ9, and Δ12 had significantly increased GFP expression. Complementation in Δ13 and Δ17 restored P_*comG*_-*gfp* expression to levels equivalent to Nef values in the biofilm (Fig. 2b).

In conclusion, these data indicate that P_*comG*_-*gfp* expression is affected by cell wall-targeting antibiotics as well as environmental cues or cellular status in biofilm where TCS13 and TCS17 play important regulatory roles.

### Biofilm growth conditions induce efficient natural transformation

Nisin and biofilm conditions were tested with tetracycline-resistant donor cells (N315Δcls2, tet^R^, or NefΔcls2, tet^R^) for their effects on transformation efficiencies. Nisin (8μg mL^−1^) had no detectable effect on transformation in Nef planktonic growth (Supplementary Fig. 6) and transformation was unaffected in the SigH-overexpressing strain (Nefh). Under biofilm-forming conditions, however, the transformation frequency in Nef increased, reaching 10^−6~7^ at day 3 (Fig. 2c), which was remarkably higher than in the planktonic growth condition (undetected, <10^−11^, n=5) (Fig. 2c). In Nefh, the transformation frequency was similar to Nef under biofilm-forming conditions in that it was higher than in the planktonic growth condition (~10^−11^, n=5) (Fig. 2c).

Figure 2d shows the time course for the transformation frequencies of Nef, Nefh, Δ13, Δ17, NefΔcomG, and NefΔcomE. Unexpectedly, we found that one of the negative control strains, the *ΔcomG* operon, was also transformable in biofilm while the *ΔcomE* operon was not (Fig. 2d). This observation is consistent with reports that the *comE* operon encodes an essential DNA incorporation channel while the *comG* operon encodes the pseudopilin that facilitates DNA access to this channel^27^. Transformation frequencies of Δ13 and Δ17 were significantly reduced at day 3 compared to Nef (Fig. 2e) but complementation restored transformation frequencies in biofilm to levels equivalent to Nef (Fig. 2e).

For experiments using planktonic competent cells, CS2 medium was indispensable for detecting transformation. We therefore evaluated if static biofilm conditions could induce natural transformation with other growth media such as TSB, BHI, RPMI, or M9 supplemented with amino acids (Supplementary Fig. 7). Natural transformation was detected in Nef and Nefh in all growth media but the efficiency was ~100 to 1000-fold lower than in CS2 medium, indicating that CS2 medium is dispensable but preferable for efficient natural transformation in Nef biofilm.

### Clinical isolates are capable of natural transformation in biofilm

As natural transformation in *S. aureus* has only been detected in N315 derivative strains that were genetically engineered to express SigH^13^, we tested the transformability of 5 unmodified clinical isolates (tetracycline susceptible) by employing the biofilm conditions described above. We found that one strain (MRSA, r59) was transformable by the tetracycline resistance marker (Supplementary Fig. 8) whereas other strains became transformation competent after introducing a SigH-expressing plasmid (pRIT-sigH), suggesting that *sigH* expression was still a limiting step in the transformation of MSSA s142 and MRSA r3. Two strains (MSSA s1567 and MRSA r408) were not transformable irrespective of the SigH-expressing plasmid. Taken together, these data reveal that the biofilm condition facilitates transformation in some, but not all, clinical isolates.

### SCC*mec* elements can be transferred in biofilm

Exploring the SCC transfer mechanism is challenging as small SCC elements can be transferred by transduction^28^, but typical staphylococcal-transducing bacteriophages are Siphoviridae with genome sizes of less than c.a. 45 kb^29^ and cannot physically accommodate an entire large SCC^30^. Conjugation can also transfer a shortened SCC*mec* II but this requires insertion of SCC into the conjugative plasmid^31^ and, to the best of our knowledge, such a plasmid-carrying SCC has not been reported in staphylococcal isolates. In this study, we propose that natural transformation in biofilms is the major mechanism for SCC transfer based on the following experimental evidence.

We tested the natural transformation of *mecA* in biofilms by using clinical isolates of MSSA as recipients and heat-killed MRSA or methicillin-resistant coagulase-negative staphylococci strains (MR-CoNS) as donors. The *mecA* transformants were selected by cefmetazole (from the cephem subgroup of the β-lactam antibiotics). We first tested 20 MSSA clinical isolates using MR-CoNS8 as donor (for 1s-20s) and other 20 MSSA using MR-CoNS3 as donor (21s-40s), and found that 6 strains were able to form colonies on cefmetazole plates. Among these 6 strains, 4 (1s, 9s, 11s, 35s) were selected for further analysis (Supplementary Table 4). These 4 MSSA strains, together with Nef and NefΔcomE as positive and negative controls, were tested for their transformability with distinct staphylococcal species and SCC types. Either *S. aureus* (including 35 transformants; 35s [CoNS17]) or MR-CoNS, along with any tested SCC*mec* (I, II, III, IVa), could serve as the donor with the detected efficiencies ranging from c.a. 10^−8^ to 10^−7^, generating up to ~ 160 colonies from a single-well biofilm containing 10^9^ c.f.u recipient cells. To confirm this observation as natural transformation, we deleted the *comE* operon of 9s and found that this mutant was non-transformable using N315 as the donor (Supplementary Table 4).

Some of these MSSA recipients (1s, 9s, 11s, but not 35s) were transformed by the chromosomal tetracycline resistance gene while no transformation was detected using the pT181 plasmid in any transformable strain (Supplementary Table 4, Nef-pT181). The question as to transfer of other plasmids by natural transformation under biofilm conditions remains unanswered.

Figure 3a shows the result of *mecA* colony PCR for multiple transformants. All transformants derived from 35s showed the *mecA* signal and the minimum inhibitory concentrations (MICs) of cephems (cefmetazole and cefoxitin) were increased in these transformants, demonstrating their conversion to MRSA (Supplementary Table 5). In contrast, some transformants showed lower intensities in *mecA* signal compared with 35s transformants (Fig. 3a). Moreover, the MIC values in these transformants were relatively lower than 35s transformants, suggesting that 35s, but not others, could stably accommodate the *mecA* gene. The stability test of cefoxitin resistance showed that the 35s transformant (35s [CoNS15]) sustained the resistance in the absence of β-lactam but 1s, 9s, and 11s derivatives tested swiftly lost their resistance (Fig. 3b). Disk diffusion test (Fig. 3c) confirmed the reduced susceptibility of the stable transformant 35s[CoNS15] to cefoxitin (CFX) and oxacillin (MPI). On the other hand, the unstable 9s[CoNS15] was categorized into the MSSA criteria, though the inhibitory zone of CFX slightly decreased and colony appeared on the edge of the inhibitory zone of MPI. Notably, we detected stable 9s transformants when N315 was used as donor. This suggests that SCC*mec* type and recipient strain are drivers of SCC*mec* stability in transformants. PCR analysis showed that the full-length transferred SCC*mec* IVa was present in the stable 35s transformants (Fig. 3d). Full-size SCC*mec* II was detected in transformants of Nef and 9s but was shortened in 35s transformants, possibly due to the elimination of mobile elements Kdp, Tn554, and IS431-pUB110 (Fig. 3e). Stability and SCC intactness in transformants are summarized in Fig. 3f. Collectively, these observations demonstrate that SCC*mec* elements can be transferred to MSSA strains by natural transformation in biofilm.

### SCC*mec* transformation depends on *ccrAB* recombinase genes and an intact *attB* site

SCC carries *ccr* genes encoding a dedicated excision and integration system but there is scarce evidence of the mechanistic requirements of this system for intercellular HGT. To determine whether SCC*mec* transformation is mediated by CcrAB (cassette chromosome recombinases) or general homologous recombination, we deleted the *ccrAB* gene from the SCC*mec* II element (N315ΔccrAB) and evaluated its ability to serve as SCC*mec* donor for 9s and Nef strains. SCC*mec* transformants could not be obtained using N315ΔccrAB as a donor (Supplementary Table 4), suggesting that the site-specific excision and integration of SCC*mec* mediated by *ccrAB* is essential for transformant generation. We also generated mutations in the *attB* sequence on the recipient side (NefattB*) and this did not generate SCC*mec* transformants when N315 was used as donor (Supplementary Table 4). This novel evidence points to the *ccrAB-attB*-dependent SCC transfer system as critical for SCC*mec* transformation.

### Bacitracin inhibits natural transformation

Our finding that cell wall-targeting antibiotics affect P_*comG*_ activity (Supplementary Fig. 4) suggests that they may also affect natural transformation. To test this point, we treated cells growing in biofilm with either bacitracin, vancomycin, or nisin for three days. Low-concentration bacitracin treatment (0.5 μg mL^−1^) reduced the transformation efficiency while 5 μg mL^−1^ completely prevented the detection of transformants in Nef (Fig. 4a). Vancomycin treatment, on the other hand, had no significant effect on natural transformation in this strain at all tested concentrations (Fig. 4b). Nisin treatment at 16 μg mL^−1^ significantly increased the transformation efficiency (Fig. 4c).

## Discussion

This study demonstrated HGT of SCC*mec* by natural transformation and provides mechanistic information on the pathway of MRSA emergence. SCC, an MGE shared among staphylococcal species and *Macrococcus caseolyticus*^32^, is responsible for dissemination of virulence factors and resistance genes such as capsule synthesis genes (SCC*cap*)^33,34^, the fusidic acid resistance gene (SCC*fus*)^35^, and the methicillin resistance gene (SCC*mec*) (see comprehensive review^36^). Since its discovery, SCC*mec* has been a research focus of extensive efforts to clarify the global emergence and dissemination of MRSA. In 1961, the first MRSA, which carried the type I SCC*mec,* was isolated in the United Kingdom. Types II and III, identified in the early 1980s in Japan and New Zealand, were clinically isolated and are reported as the largest types among SCCs^37,38^. SCC*mec* IV and V were described in United States and Australia but are relatively small and found primarily in community-acquired MRSAs^39,40^. While Types I to V are dominant and widely distributed, diverse new variants have been reported (Types VI – XIII). The origins of SCC*mec* are unclear but ancestral forms have been identified in coagulase-negative staphylococci such as *S. sciuri, S. fleuretti, S. xylosus, S. hominis,* and *M. caseolyticus*^32,41,42^. Ccr recombinases were found to mediate the excision and insertion of SCC at the *attB* locus (*attL*/*attR* after SCC*mec* integration)^10^, with CcrA and CcrB for SCC type I ~type V and CcrC for type V. These *ccrAB* genes are expressed in minor subpopulations and the excised circular SCC is thought to serve as a donor for horizontal transmission^43^. Both proteins are required for the proper excision of SCC*mec* from the chromosome and its integration into the *attB* site after transduction as a part of the artificial plasmid^44^.

Despite such well-established epidemiological evidence of interspecies SCC*mec* movement and *ccr*-dependent excision/insertion, the major intercellular transmission mechanism has remained enigmatic for half a century. Transfer of the methicillin resistance gene was first demonstrated by transduction^45^ and by pseudo-competence^46^ which, at that time was described as ‘transformation’ but, after discovery of the phage component, is now termed ‘pseudo-competence’ or ‘pseudo-transformation’. Transduction has been suggested as a preferable transfer route for some types of SCC*mec* (albeit with a < 45 kb capacity limit of the bacteriophage capsid) and SCC*mec* fragments are detectable in bacteriophage capsids^47,48^, making the transfer of short SCC*mec* (types IVa and I) observable by transduction among compatible strains^28^. However, major deletions are occasionally associated with transduction^28^ and successful integration into the recipient chromosome requires homologous flanking sequences, suggesting that this transduction relies on homologous recombination rather than the *ccr*-mediated system. Conjugation has also been suggested as a possible mechanism for SCC*mec* transfer^31^. However, successful transfer requires donor manipulation by overexpressing the *ccr* recombinase to capture a shortened SCC*mec* into a conjugative plasmid^31^ while spontaneous and large element transfers have not been demonstrated.

The SCC transformation observed in this study was dependent on CcrAB-mediated excision/integration. To the best of our knowledge, this study is the first to show that natural intracellular SCC HGT requires the *ccr-attB* system and, based on this evidence, we propose that natural transformation is the major route for SCC*mec* transmission. The high SCC transformation efficiency, up to 10^−7^ (Supplementary Table 4), strongly supports the historical, independent transfers of distinct SCC types to *S. aureus*. Although transduction remains as a candidate HGT mechanism for short SCC*mec,* the extent of *ccr* involvement in this process remains elusive.

It has been suggested that *mecA* acquisition and expression in *S. aureus* is a fitness cost and the process of obtaining β-lactam resistance is complex, involving multiple mutations and metabolic adaptations^49,50,51^. It was previously suggested that different *S. aureus* genetic backgrounds offer different capacities to accommodate *mecA*^11,52^ and we observed a similar effect as two of our tested MSSA strains (9s and 35s) could stably accommodate the transferred *mecA* whereas 1s and 11s could not (Fig. 3). The methodology of SCC*mec* transfer established in this study would be invaluable to detail the genetic factors that define *mecA* stability.

The present study clarified that biofilm conditions are preferable for natural transformation in *S. aureus*. All tested factors that positively affect biofilm formation also increased transformation efficiency, such as CS2 medium, static growth (Fig. 2), TCS13, and TCS17 (Fig. 2), but how the biofilm structure increases transformation efficiency remains unknown. It is likely that transformation following competence gene expression is facilitated in biofilm as the P_*comG*_ reporter expression was reduced but transformation frequency was increased in biofilm compared to the planktonic state (Figs. 1,2). In addition, the transformation efficiency of SigH-overexpressing cells was higher in biofilm than in planktonic growth (Fig. 2c). In order to detect transformation in biofilm, however, it was crucial to use heat-killed donor cells rather than purified chromosomal DNA. This might be consistent with the fact that nuclease production is a common characteristic across all strains of *S. aureus* and also occurs in biofilms^53^. Alternatively, it is possible that experimentally added, purified DNA cannot serve as a transformation donor since extracellular DNA is known to be tightly sequestered in biofilm^54,55^. It is interesting to note that, in *Streptococcus pneumoniae*, non-competent cells undergo lysis by bacteriocins and fratricins released by neighboring competent cells^56^ but the presence of such a dedicated mechanism for DNA supply is not known in the *Staphylococcus* genus.

Staphylococcal infections are intimately associated with biofilm formation^57^ as it provides protection against antimicrobial treatment and host clearance mechanisms^58^ while contributing to the prolonged infection and colonization that facilitates the dissemination of drug-resistant strains^59^. Our finding that *S. aureus* can develop natural transformation in biofilm conditions emphasizes the additional role of biofilms in promoting HGT as well as transduction and conjugation^1,60^. Additionally, cells release phage at higher frequency than in planktonic conditions^61^ and subsequent cell lysis in biofilm would create an ample supply of genetic material for non-lysed cells. Interestingly, expressing SigH was shown to stabilize phage lysogeny^62^, implicating a co-evolution of distinct HGT mechanisms in staphylococcal biofilms. Mixed biofilms of *S. aureus* and other staphylococci formed during commensal state or co-infections are thus general hot spots for HGT.

Crucially, natural transformation can transfer longer DNA fragments, such as SCC*mec* II (Fig. 3e), that are too large to be packed into the typical staphylococcal bacteriophages^13^. Natural transformation cannot be abolished by inactivating the donor, unlike other HGT mechanisms such as phage transduction, conjugation, and the staphylococcal pathogenicity island-helper phage system. In order to counter staphylococcal evolution by SCC systems, specific control methods against transformation are therefore necessary and the recipient cell’s signal transduction systems (*e.g.*, TCS17) might serve as a promising target. The finding of the inhibitory effect of bacitracin (Fig. 4a) may also serve as an attractive future direction for experimental studies.

## Methods

### Bacterial strains and culture conditions

Bacterial strains and plasmids used in this study are listed in Supplementary Table 1. Clinical staphylococcal samples (40 MSSA isolates and 7 MR-CoNS isolates) were collected from the Kanto area of Japan. Unless otherwise indicated, staphylococci were grown in Trypticase Soy Broth (TSB). *E. coli* strains were grown in LB. Cultures were incubated at 37°C either with shaking (180 r.p.m) or statically. Where required for selection, culture medium was supplemented with chloramphenicol (12.5 μg mL^−1^), kanamycin (100 μg mL^−1^), tetracycline (5 μg mL^−1^), cefoxitin (4 μg mL^−1^), cefmetazole (4 μg mL^−1^), or ampicillin (100 μg mL^−1^).

### Construction of deletion and substitution mutants

Each mutant was constructed from Nef or 9s by double-crossover homologous recombination using the pMADtet vector^13^ (Supplementary Table 1). Briefly, two fragments flanking the upstream (primers A, and B, Supplementary Table 2) and downstream (primers C, and D, Supplementary Table 2) regions of the locus targeted for deletion (or substitution) were amplified by PCR. The PCR products (AB and CD fragments) were used as template to generate the construct AD by overlapping PCR, using the primers A and D depending on locus (Supplementary Table 2). Product AD was cloned into the *Bam*H I – *Sal* I site of pMADtet to generate the vectors for TCS deletions (pMADtet-Δ3 to Δ17), pMADtet-ΔccrAB, and pMADtet-attB* (Supplementary Table 1). In terms of *attB* substitution, primers B and C were designed not to change the coding amino acid sequence of OrfX. The plasmids, purified from *E.coli* DH5α, were introduced into Nef, after passaging through RN4220. Mutants (tetracycline sensitive, β-galactosidase negative) were selected as described previously^13,63^ and the absence of the target gene was confirmed by PCR using the primers E and F (Supplementary Table 2). The *attB* substitution mutant was confirmed by restriction digestion (*Hin*dIII: included in the designed primers attB-B and attB-C) of the PCR product generated by primers E and F (Supplementary Table 2). NefΔcls2 strain was created by transduction using the donor N315Δcls2 (carries a tetracycline resistance gene at the *cls2* locus^64^) (Supplementary Table 1).

### Complementation of ΔTCSs

For *in trans* complementation, each TCS gene (including its own promoter) was amplified by PCR using chromosomal DNA from Nef as genomic template. The PCR product was cloned into the *Eco*R I – *Bam*H I site of pHY300PLK (Takara) to generate the complementation plasmids pHY-12, pHY-13, and pHY-17 (Supplementary Table 1). These plasmids were introduced into the corresponding mutants after passaging through RN4220.

For chromosomal complementation of Δ13 and Δ17, each TCS its flanking region were amplified by PCR using primers G and H (Supplementary Table 2). Product was cloned into *Bam*H I – *Sal* I site of pMADtet to generate the vectors pMADtet-13 and pMADtet-17 (Supplementary Table 1). The plasmids were purified from *E.coli* DH5α and introduced into the corresponding ΔTCS after passaging through RN4220. Complemented mutants (tetracycline sensitive, β-galactosidase negative) were selected as described previously^13,63^ and the presence of the restored gene was confirmed by PCR using the primers E and F (Supplementary Table 2).

### Antimicrobial susceptibility testing

MIC assays were conducted in a 96-well microtiter plate (round bottom). Overnight bacterial cultures were diluted 1:2000 in appropriate medium and 100 μL aliquots were used to inoculate wells containing TSB or CS2 media supplemented with twofold serial dilutions of antibiotics (vancomycin, bacitracin, nisin, cefmetazole, or cefoxitin). The plates were statically incubated for 20 h at 37 °C. The MIC was determined by the lowest concentration of antibiotic at which growth was inhibited.

Disk diffusion testing was conducted according to CLSI standard using direct colony suspension method. Briefly, glycerol stocks of 9s, 35s, CoNS15, and 35s[CoNS15] were streaked on drug-free TSA. Unstable transformant (9s[CoNS15]) was streaked on TSA supplemented with 4 μg mL^−1^ cefoxitin. Emerged colonies were suspended in 0.85% NaCl and turbidity was adjusted to 0.5 McFarland standard. The inocula were swabbed on Mueller-Hinton agar and the antibiotic disks of oxacillin (1 μg), and cefoxitin (30 μg) (KB disks, Eiken Chemical) were used for susceptibility testing. Zone of inhibition was determined following 18 h of incubation at 35 °C.

### Measurement of *comG* promoter activity by a GFP reporter assay

The reporter plasmid of P_*comG*_-gfp (pMK3-com-gfp)^13^ was introduced into each strain by electroporation after passaging through RN4220. Reporter strains were grown overnight with 100 μg mL^−1^ kanamycin and diluted 1:200 in the appropriate medium supplemented with vancomycin, bacitracin, or nisin as appropriate. Next, 200 μl of these diluted cultures were placed in a transparent, 96-well flat-bottomed microplate (Thermo Scientific, MA, USA) before continuous incubation (with shaking) at 37°C in a multimode plate reader (2300 Enspire™, PerkinElmer^®^). Changes in the fluorescence intensity and OD_600_ were measured over 36 hours with 30 min intervals. The fluorescence intensity was normalized by the OD_600_ value.

To count the numbers of GFP-expressing cells in planktonic culture, 50 μl of overnight culture for each reporter strain was inoculated into 10 mL of CS2 medium in a glass vial. These cells were grown at 37°C with shaking for the appropriate time period before 5 μL of the culture was placed on slide, sealed with a cover glass and observed by the fluorescence microscope (BZ-X710, Keyence). To count GFP-expressing cells grown in the static biofilm condition, bacteria in the biofilm were collected by extensive pipetting, washed and suspended in PBS, and stained by propidium iodide (40 μM final concentration) (WAKO) to distinguish dead cells (red fluorescence) from living cells. Stained bacteria were observed by the fluorescence microscope. The percentage of GFP-expressing cells was calculated by dividing the number of GFP-expressing cells by the total number of living cells.

### Natural transformation assay

Donors used in natural transformation assays either include a purified plasmid (pHY-300PLK) (Supplementary Fig. 6) or heat-killed cells (Figs. 2c,d,e, 3, 4, Supplementary Figs. 6,7,8, Supplementary Table 4). Heat-killed donor was prepared by diluting an overnight culture 1:20 in TSB and growing with shaking for 3 h at 37°C. Next, cells were harvested and suspended in 5 ml PBS before heating in boiling water for 10 min. The absence of viable cells was confirmed by plating on TSB agar.

Natural transformation assays in planktonic condition (Fig. 2c, Supplementary Fig. 6) were carried out as previously described^13^ with some modifications. Briefly, 500 μl of recipient cells from overnight cultures were washed and inoculated in 10 ml CS2, supplemented with 8 μg mL^−1^ nisin when required, and were grown for 8 h at 37°C with shaking. Cells were then harvested and suspended in fresh 10 ml CS2 containing 10 μg of purified pHY-300PLK plasmid (Supplementary Fig. 6) or 5×10^10^ heat-killed N315Δcls2 cells (Fig. 2c). Growth was continued for additional 2 h (Supplementary Fig. 6) or 3 days (Fig. 2c) before pouring into melted BHI agar supplemented with 5 μg mL^−1^ tetracycline to select for transformants.

To detect natural transformation in biofilms (Fig. 2, 3, 4, Supplementary Figs. 7,8, Supplementary Table 4), overnight cultures of recipient cells were diluted 1:200 in TSB and grown with shaking for 3 h at 37°C before harvesting 750 μl of the culture, washing it and suspending it in appropriate growth medium (CS2 medium, TSB, BHI, RPMI 1640, or M9 supplemented with amino acids as in CS2 medium^13^). The 750 μl of cell suspension (10^8^ c.f.u mL^−1^) was distributed in a 6-well flat bottom polystyrene plate (Costar^®^, Corning) and 250 μl of heat-killed donor cells (10^9^ c.f.u mL^−1^) were added. The total volume of growth medium was adjusted to 1.5 mL per well. The 6-well plate was statically incubated at 37°C and medium was refreshed every 24 h. After incubation for an appropriate time period, the biofilm was harvested by extensive pipetting and was poured into BHI agar supplemented with 5 μg mL^−1^ tetracycline or 4 μg mL^−1^ cefmetazole, depending on the donor used. Unless otherwise indicated, all transformation frequencies from the assays were determined after 3 days of growth in biofilm. Generated colonies in cefmetazole plates were replicated onto fresh agar plates containing 4 μg mL^−1^ cefmetazole to confirm the acquired resistance, and those that could grow were counted as transformants. Transformation frequency was calculated as the ratio of the number of transformants to the total c.f.u. Non-detected values were assigned half the value of the detection limit of the strain for the calculation of mean values (Figs. 2, 4) and statistical analyses.

### Stability test for cefoxitin resistance

Glycerol stocks of *mecA* transformants were streaked on BHI plates supplemented with 4 μg mL^−1^ cefoxitin. Emerged single colonies were inoculated into BHI medium supplemented with 4 μg mL^−1^ cefoxitin and grown with shaking at 37 °C for 12 h. This culture was then diluted 1:1000 in drug-free BHI medium and grown for another 12 h. After 12 h cells were plated on drug-free BHI agar and were replicated on BHI agar supplemented with 4 μg mL^−1^ cefoxitin to assess the population percentage that maintained growth ability under selective pressure from the β-lactam drug.

### Biofilm staining and quantification

Biofilm formation was assessed in 96-well plates^65^. Overnight cultures were diluted 1:200 in CS2 medium and 200 μl were transferred to each well. Following 24 h of static incubation at 37 °C, the nonadherent cells in medium were aspirated and the wells were stained with 200 μl of 0.1% crystal violet for 15 min. The wells were then gently washed 3 times with 200 μl of PBS to remove residual stain before air drying. For biomass quantification, 100 μl of 96% ethanol was added to the wells and incubated at room temperature for 10 min to solubilize the stain. The absorbance at 595 nm of the resolved stain was measured by plate reader.

To assess biofilm development in the natural transformation assay, the same cell suspension was inoculated in a 6-well plate containing CS2 medium (1.5 mL total volume per well). These plates were statically incubated for 96 h at 37°C. Every 24 h, formed biofilms were stained and quantified. For biofilm staining, the nonadherent cells were removed and the wells were stained with 1.5 mL of 0.1% crystal violet for 15 min. Excess stain was removed by distilled water and the wells were air dried. For biomass quantification, 750 μl of 96% ethanol was added to the wells for 10 min at room temperature and the absorbance at 595 nm of the resolved stain was measured. Cell-free wells were used as blanks.

### SCC*mec* typing and long amplification by PCR

SCC*mec* typing for MR-CoNS donors was performed by multiplex PCR as previously described^66^. Long amplifications of SCC*mec* were performed using KOD One PCR master mix (TOYOBO) according to the manufacturer’s instructions.

### Statistical analysis

Statistical analyses were performed by GraphPad Prism (GraphPad Software, version 8.4.3). The differences between groups were analysed by either t-test or one-way ANOVA followed by Dunnett’s or Tukey’s multiple comparisons test as indicated in figure legends. The log values of natural transformation frequencies were analysed statistically. P values below 0.05 were considered statistically significant.

## Acknowledgement

This study was supported by Takeda Science Foundation, Pfizer Academic Contributions, JSPS KAKENHI Grant Number 25860313 and 18H02652 (to KM), Program to Disseminate Tenure Tracking System, MEXT (to RLO). We would like to express our appreciation to Dr. Teruyo Ito for valuable discussion, and Ms. Yoshimi Tsutsumi, Mr. Tin Ming Tan, Mr. Bobby Sookhoo, and Ms. Clara Effenberger for their experimental help. We would also like to thank Dr. Bryan J. Mathis, Medical English Communications Center, University of Tsukuba, for language revision of this manuscript.

## Supplementary materials

**Supplementary Table 1. Bacterial strains and plasmids used in this study.**

**Supplementary Table 2. List of primers used in this study**

**Supplementary Table 3. MIC (μg mL^−1^) of cell wall-targeting antibiotics in CS2 medium or in TSB.**

**Supplementary Table 4. Intra-and interspecies transformation of distinct SCC***mec* **elements under biofilm growth conditions.**

**Supplementary Table 5. MIC (μg mL^−1^) of cefmetazole and cefoxitin in TSB.**

**Supplementary Figure 1. Reporter assay for***comG* **promoter activity.** Nef carrying the P_*comG*_-*gfp* reporter was grown in either CS2 medium (a) or TSB (b) with shaking. Fluorescence intensity (green line) and OD_600_ (blue line) were measured every 30 min. The mean of n = 3 independent experiments is shown. Error bars represent s.d.

**Supplementary Figure 2. Growth curves of Nef and its derivative ΔTCS.** Cells were grown in either TSB (**a**) or CS2 medium (**b**) with shaking. OD_600_ was measured every 30 min. The mean of n = 3 independent experiments is shown. Error bars are omitted for clarity.

**Supplementary Figure 3. Biofilm formation is impaired in the TCS13 and TCS17 mutants.**

**(a)** Nef and its derivatives were statically grown in CS2 medium in a 96-well plate for 24 h.

**(b)** Time-course of biofilm development in Nef and its derivatives. The cells were statically grown in CS2 medium in a 6-well plate for up to 4 days. (**b, middle**) The biofilms were stained with crystal violet and the absorbance at 595 nm was measured. The mean of n = 2-3 independent experiments is shown. Error bars represent SD. (**b, right**) CFU of the cells after 24 h in the biofilm.

**Supplementary Figure 4.** *comG* **promoter activity is affected by cell wall-targeting antibiotics.** Nef-GFP was treated with subinhibitory concentrations of vancomycin (**a**), bacitracin (**b**), or nisin (**c**). Cells were grown in CS2 medium with shaking for 24 h. Fluorescence (RFU) and OD_600_ were measured every 30 min. Data shown are either relative RFU/OD_600_ values (**left panels**) or increases in RFU/OD_600_ values during 8-24 h of growth (**right panels**).

**Supplementary Figure 5. *comG* promoter activity is affected by multiple TCSs in biofilm.** Nef and its derivative ΔTCSs carrying the P_*comG*_-*gfp* reporter were statically grown in CS2 medium in biofilm growth conditions for 3 days. The percentage of GFP-expressing cells was calculated as observed by fluorescence microscopy. The mean of n = 3-7 independent experiments is shown. Error bars represent s.d. Statistical significance was determined by one-way ANOVA with Tukey’s multiple comparison test. *P<0.05, ****P<0.0001.

**Supplementary Figure 6. Nisin does not induce natural transformation in Nef**. Nef and its derivatives were grown in CS2 medium with or without nisin (8 μg mL^−1^). Transformation frequencies were determined after 10 h of planktonic growth. The transformants were selected by tetracycline. The mean of two independent experiments is shown with s.d. ND < 10^−9^.

**Supplementary Figure 7. Nutrient-poor culture media are preferable for natural transformation under biofilm growth conditions**. Nef, Nef-h, and Nef-ΔcomE were statically grown in different growth media, including CS2, TSB, BHI, RPMI, and M9 supplemented with amino acids (M9+aa). Transformation frequencies were determined at day 3. Dotted lines represent the detection limit of the strains in each growth medium. The mean and s.d. are shown. Circles represent independent experiments.

**Supplementary Figure 8. Transformation frequencies of clinical isolates.** Wild-type (WT) *S. aureus* and its derivative carrying the SigH expression plasmid pRIT-sigH (WT-h) were tested for transformation after 2 days of static growth in CS2 medium. The transformants were selected by tetracycline. The mean of n = 2-3 independent experiments is shown. Error bars represent s.d.

## Notes

### Competing Interest Statement

The authors have declared no competing interest.

