## supplementary materials for "MRSA emerges through natural transformation"

**Supplementary Table 1. Baterial strains and plasmids used in this study**

| Strain or plasmid | Description | Source |
|---|---|---|
| **Strains** | | |
| N315 | Pre-MRSA, SCC*mec* II, Km$^R$, Erm$^R$ | 88 |
| N315ΔccrAB | N315 lacking *ccrAB* locus | This study |
| N315ex | N315 cured of SCC*mec*, Km$^S$, Erm$^R$ | 37 |
| Nef | N315ex w/oφ (N315 cured of SCC*mec* and the φN315 prophage) | 13 |
| Nefh | Nef carrying pRIT-sigH | 13 |
| NefΔcomG | Nef lacking *comG* operon | 13 |
| NefΔcomE | Nef lacking *comE* operon | 13 |
| Δ3 | Nef lacking TCS3 | This study |
| Δ4 | Nef lacking TCS4 | This study |
| Δ5 | Nef lacking TCS5 | This study |
| Δ6 | Nef lacking TCS6 | This study |
| Δ7 | Nef lacking TCS7 | This study |
| Δ8 | Nef lacking TCS8 | This study |
| Δ9 | Nef lacking TCS9 | This study |
| Δ10 | Nef lacking TCS10 | This study |
| Δ11 | Nef lacking TCS11 | This study |
| Δ12 | Nef lacking TCS12 | This study |
| Δ13 | Nef lacking TCS13 | This study |
| Δ14 | Nef lacking TCS14 | This study |
| Δ15 | Nef lacking TCS15 | This study |
| Δ16 | Nef lacking TCS16 | This study |
| Δ17 | Nef lacking TCS17 | This study |
| Δ13 (13) | Nef ΔTCS3 mutant with chromosomal complementation | This study |
| Δ17 (17) | Nef ΔTCS17 mutant with chromosomal complementation | This study |
| Nef-GFP | Nef carrying pMK3-com-gfp | This study |
| Δ3-GFP | Nef ΔTCS3 mutant carrying pMK3-com-gfp | This study |
| Δ4-GFP | Nef ΔTCS4 mutant carrying pMK3-com-gfp | This study |
| Δ5-GFP | Nef ΔTCS5 mutant carrying pMK3-com-gfp | This study |
| Δ6-GFP | Nef ΔTCS6 mutant carrying pMK3-com-gfp | This study |
| Δ7-GFP | Nef ΔTCS7 mutant carrying pMK3-com-gfp | This study |
| Δ8-GFP | Nef ΔTCS8 mutant carrying pMK3-com-gfp | This study |
| Δ9-GFP | Nef ΔTCS9 mutant carrying pMK3-com-gfp | This study |
| Δ10-GFP | Nef ΔTCS10 mutant carrying pMK3-com-gfp | This study |
| Δ11-GFP | Nef ΔTCS11 mutant carrying pMK3-com-gfp | This study |
| Δ12-GFP | Nef ΔTCS12 mutant carrying pMK3-com-gfp | This study |
| Δ13-GFP | Nef ΔTCS13 mutant carrying pMK3-com-gfp | This study |
| Δ14-GFP | Nef ΔTCS14 mutant carrying pMK3-com-gfp | This study |
| Δ15-GFP | Nef ΔTCS15 mutant carrying pMK3-com-gfp | This study |
| Δ16-GFP | Nef ΔTCS16 mutant carrying pMK3-com-gfp | This study |
| Δ17-GFP | Nef ΔTCS17 mutant carrying pMK3-com-gfp | This study |
| Δ12-GFP (pHY) | Nef ΔTCS12 mutant carrying pMK3-com-gfp and pYH300PLK (empty vector) | This study |
| Δ12-GFP (pHY-12) | Nef ΔTCS12 mutant carrying pMK3-com-gfp and pYH-12 (TCS12 complementary strain) | This study |
| Δ13-GFP (pHY) | Nef ΔTCS13 carrying pMK3-com-gfp and pYH300PLK (empty vector) | This study |
| Δ13-GFP (pHY-13) | Nef ΔTCS13 mutant carrying pMK3-com-gfp and pYH-13 (TCS13 complementary strain) | This study |
| Δ17-GFP (pHY) | Nef ΔTCS13 carrying pMK3-com-gfp and pYH300PLK (empty vector) | This study |
| Δ17-GFP (pHY-17) | Nef ΔTCS13 mutant carrying pMK3-com-gfp and pYH-17 (TCS17 complementary strain) | This study |
| Δ13-GFP (13) | Nef ΔTCS3 mutant carrying pMK3-com-gfp with chromosomal complementation | This study |
| Δ17-GFP (17) | Nef ΔTCS17mutant carrying pMK3-com-gfp with chromosomal complementation | This study |
| N315Δcls2-tet$^R$ | N315Δcls2 mutant, Tet$^R$ | Tsai, 2011 |
| NefΔcls2-tet$^R$ | NefΔcls2 mutant, Tet$^R$ (transduction of Tet$^R$ from N315Δcls2) | This study |
| NefattB* | Nef with mutated *attB* site | This study |
| Nef-pT181 | Nef carrying pT181 | This study |
| s142 | Clinical isolate, MSSA | 89 |
| s142-h | s142 carrying pRIT-sigH | This study |
| s1567 | Clinical isolate, MSSA, clonal complex 5 | 89 |
| s1567-h | s1567 carrying pRIT-sigH | This study |
| r3 | Clinical isolate, MRSA, SCC*mec*II, clonal complex 5 | 89 |
| r3-h | r3 carrying pRIT-sigH | This study |
| r59 | Clinical isolate, MRSA, SCC*mec*II, clonal complex 5 | 89 |
| r59-h | r59 carrying pRIT-sigH | This study |
| r408 | Clinical isolate, MRSA, SCC*mec*II, clonal complex 5 | 89 |
| r408-h | r408 carrying pRIT-sigH | This study |
| RN4220 | Derivative of 8325-4, restriction minus, modification plus | 90 |
| DH5α | *E. coli* competent cell | Toyobo Ltd., Japan |
| 1s | MSSA clinical isolate | This study |
| 9s | MSSA clinical isolate | This study |
| 9sΔcomE | 9s mutant lacking *comE* operon | This study |
| 11s | MSSA clinical isolate | This study |
| 35s | MSSA clinical isolate | This study |
| MR-CoNS3 | Methicillin resistant CoNS3, *S. epidermidis*, SCC*mec* nontypeable (isolated from human blood) | This study |
| MR-CoNS8 | Methicillin resistant CoNS8, *S. lugdunensis*, SCC*mec* nontypeable (isolated from human blood) | This study |
| MR-CoNS9 | Methicillin resistant CoNS9, *S. epidermidis*, SCC*mec* III (isolated from human blood) | This study |

| | | |
|---|---|---|
| MR-CoNS10 | Methicillin resistant CoNS10, *S. epidermidis*, SCC*mec* IVa (isolated from human blood) | This study |
| MR-CoNS11 | Methicillin resistant CoNS11, *S. epidermidis*, SCC*mec* IVa / I (isolated from human blood) | This study |
| MR-CoNS15 | Methicillin resistant CoNS15, *S. epidermidis*, SCC*mec* IVa (isolated from human blood) | This study |
| MR-CoNS16 | Methicillin resistant CoNS16, *S. caprae*, SCC*mec* I (isolated from human blood) | This study |
| MR-CoNS17 | Methicillin resistant CoNS17, *S. epidermidis*, SCC*mec* IVa (isolated from human blood) | This study |
| **Plasmids** | | |
| pMADtet | pMAD derivative, $Amp^R$ (*E. coli*), $Erm^R$, $Tet^R$ (*S. aureus*) | 13 |
| pMADtet-Δ3 | Vector for deletion of TCS3 locus, $Amp^R$ (*E. coli*), $Erm^R$, $Tet^R$ (*S. aureus*) | This study |
| pMADtet-Δ4 | Vector for deletion of TCS4 locus, $Amp^R$ (*E. coli*), $Erm^R$, $Tet^R$ (*S. aureus*) | This study |
| pMADtet-Δ5 | Vector for deletion of TCS5 locus, $Amp^R$ (*E. coli*), $Erm^R$, $Tet^R$ (*S. aureus*) | This study |
| pMADtet-Δ6 | Vector for deletion of TCS6 locus, $Amp^R$ (*E. coli*), $Erm^R$, $Tet^R$ (*S. aureus*) | This study |
| pMADtet-Δ7 | Vector for deletion of TCS7 locus, $Amp^R$ (*E. coli*), $Erm^R$, $Tet^R$ (*S. aureus*) | This study |
| pMADtet-Δ8 | Vector for deletion of TCS8 locus, $Amp^R$ (*E. coli*), $Erm^R$, $Tet^R$ (*S. aureus*) | This study |
| pMADtet-Δ9 | Vector for deletion of TCS9 locus, $Amp^R$ (*E. coli*), $Erm^R$, $Tet^R$ (*S. aureus*) | This study |
| pMADtet-Δ10 | Vector for deletion of TCS10 locus, $Amp^R$ (*E. coli*), $Erm^R$, $Tet^R$ (*S. aureus*) | This study |
| pMADtet-Δ11 | Vector for deletion of TCS11 locus, $Amp^R$ (*E. coli*), $Erm^R$, $Tet^R$ (*S. aureus*) | This study |
| pMADtet-Δ12 | Vector for deletion of TCS12 locus, $Amp^R$ (*E. coli*), $Erm^R$, $Tet^R$ (*S. aureus*) | This study |
| pMADtet-Δ13 | Vector for deletion of TCS13 locus, $Amp^R$ (*E. coli*), $Erm^R$, $Tet^R$ (*S. aureus*) | This study |
| pMADtet-Δ14 | Vector for deletion of TCS14 locus, $Amp^R$ (*E. coli*), $Erm^R$, $Tet^R$ (*S. aureus*) | This study |
| pMADtet-Δ15 | Vector for deletion of TCS15 locus, $Amp^R$ (*E. coli*), $Erm^R$, $Tet^R$ (*S. aureus*) | This study |
| pMADtet-Δ16 | Vector for deletion of TCS16 locus, $Amp^R$ (*E. coli*), $Erm^R$, $Tet^R$ (*S. aureus*) | This study |
| pMADtet-Δ17 | Vector for deletion of TCS17 locus, $Amp^R$ (*E. coli*), $Erm^R$, $Tet^R$ (*S. aureus*) | This study |
| pMADtet-ΔccrAB | Vector for deletion of ccrAB locus, $Amp^R$ (*E. coli*), $Erm^R$, $Tet^R$ (*S. aureus*) | This study |
| pMADtetcomEII | Vector for deletion of comE locus, $Amp^R$ (*E. coli*), $Erm^R$, $Tet^R$ (*S. aureus*) | 13 |
| pMADtet-13 | Vector for complementation of TCS13 locus, $Amp^R$ (*E. coli*), $Erm^R$, $Tet^R$ (*S. aureus*) | This study |
| pMADtet-17 | Vector for complementation of TCS17 locus, $Amp^R$ (E. coli), $Erm^R$, $Tet^R$ (S. aureus) | This study |
| pMADtet-attB* | Vector for mutational substitution of attB locus, $Amp^R$ (*E. coli*), $Erm^R$, $Tet^R$ (*S. aureus*) | This study |
| pMK3-com-gfp | $P_{comG}$-*gfp* transcriptional fusion in pMK3, $Amp^R$ (*E. coli*), $Tet^R$ *(S. aureus)* | 13 |
| pRIT-sigH | *sigH* overexpressing plasmid, $P_{spa}$, SD sequence modified into SigA type sequence | 13 |
| pHY300PLK | *Shuttle vector, ori-pAMα1, AmpR (E.coli), TetR (S. aureus)* | Takara, Japan |
| pHY-12 | TCS12 complementation plasimd, pHY300PLK-TCS12 | This study |
| pHY-13 | TCS13 complementation plasimd, pHY300PLK-TCS13 | This study |
| pHY-17 | TCS17 complementation plasimd, pHY300PLK-TCS17 | This study |

**Supplementary Table 2 Baterial strains and plasmids used in this study**

| Primer | Sequence | Reference |
|---|---|---|
| **ΔTCS construction (pMADtet targetting vector construction)** | | |
| TCS3-A | GGAGGATCCAGAAGATTTTGTTCCCTATC | This study |
| TCS3-B | CATTGAATCATCTCCAAAAATTTATGATG | This study |
| TCS3-C | TTTTTGGAGATGATTCAATGAAATAGATTAGCACATAACTAATGATTATG | This study |
| TCS3-D | GGAGTCGACGTGTATCATTGACTGGTTTTG | This study |
| TCS4-A | GGAGGATCCGGCATGTTAGAGCATATGC | This study |
| TCS4-B | CACGATAGCACCTCAGTGAATT | This study |
| TCS4-C | GAGGTGCTATCGTGCTTTAACAGTAATCCTTTTTTTTATGCATTTTAC | This study |
| TCS4-D | GGAGTCGACACAAATAGCGGTGCGAATAAG | This study |
| TCS5-A | GGAGGATCCGAAACAATTGAACGCCTAGG | This study |
| TCS5-B | CATCCATATCACCCAATATCATTTAG | This study |
| TCS5-C | GATATTGGGTGATATGGATGTAAACATGCGTTTTGTTACTTAGAATTG | This study |
| TCS5-D | GGAGTCGACTTTGTCCACCAGACAATTCAG | This study |
| TCS6-A | GGAGGATCCAGCTCAAACACTTCCTGTTC | This study |
| TCS6-B | CATCTATTTTTTCACCTCTGTTCTTACG | This study |
| TCS6-C | GAGGTGAAAAAATAGATGTCATAATCCGATTTATTTATAAAATAAAATGC | This study |
| TCS6-D | GGAGTCGACTTTTAAGCCAAAGAGCTG | This study |
| TCS7-A | GGAGGATCCATTGCCGGAAGATGTTAAACC | This study |
| TCS7-B | CATATTTTATTCCGCCCTTTTAAAATGAC | This study |
| TCS7-C | GGGCGGAATAAAATATGATCTAAATACAAACAAAAAAGTATTGAGTG | This study |
| TCS7-D | GGAGTCGACGCAATAAATGTATCACCATCC | This study |
| TCS8-A | GGAGGATCCCGCATGATTACGCTATTTTAG | This study |
| TCS8-B | CATTTGTACACCTCATATTACGACTTTTTC | This study |
| TCS8-C | GTAATATGAGGTGTACAAATGTTTTAATCATGTCTGAGACGTCAATC | This study |
| TCS8-D | GGAGTCGACACCTTCCAAAGTCTGTTGATC | This study |
| TCS9-A | GGAGGATCCGATGATAGAGGACGTACAG | This study |
| TCS9-B | CAGGTCATACCTCCCACAC | This study |
| TCS9-C | GAGGTATGACCTGTATGGAATAAAACTGAATATAGTTATTTAAGAACGC | This study |
| TCS9-D | GGAGTCGACGTCATTACTGCGCATTGATC | This study |
| TCS10-A | GGAGGATCCGCAACACATGGTACAGCTC | This study |
| TCS10-B | CATGGTATGCCTCCCTAACTTATAA | This study |
| TCS10-C | GGAGGCATACCATGGAATAAAAATTAAAGTGGTAACAGCGC | This study |
| TCS10-D | GGAGTCGACCATTGTAAATCATCGTGCATG | This study |
| TCS11-A | GGAGGATCCGTCTTCTAACATGCGATCAAG | This study |
| TCS11-B | CATCAAATCGCTCCAATTCATTTTTAC | This study |
| TCS11-C | AATTGGAGCGATTTGATGATTTAGAATGAGCTTTTAAATATTTGTCG | This study |
| TCS11-D | GGAGTCGACTGTACCCATTACGAGTCTC | This study |
| TCS12-A | GGAGGATCCTCGTTCTATTATTGGGATGTG | This study |
| TCS12-B | CATCGATAAATCACCTCTACG | This study |
| TCS12-C | GAGGTGATTTATCGATGCAATAGTTCGTATCGAATTAAGAAAAG | This study |
| TCS12-D | GGAGTCGACACTTGGATTTGACGAACAAG | This study |
| TCS13-A | GGAGGATCCGCCACGTATCTTCAAAAGAG | This study |
| TCS13-B | CAATACGGCTCTACTTCCATAG | This study |
| TCS13-C | GAAGTAGAGCCGTATTGATATAATAAGATAATAAAGTCAGTTAACGGC | This study |
| TCS13-D | GGAGTCGACATGGTGCTGCCGTATATTTG | This study |
| TCS14-A | GGAGGATCCTGATTCACCGTTATAGTGTTG | This study |
| TCS14-B | CATAACCTTCACCTCGATAGC | This study |
| TCS14-C | CGAGGTGAAGGTTATGAAATAATTAAATAAAAAAGATCGCTGCC | This study |
| TCS14-D | GGAGTCGACGTCTGTGAGTAAAGGTGTATG | This study |
| TCS15-A | GGAGGATCCTATGCGCATACATTGTGTCG | This study |
| TCS15-B | CATAGCTATAAACTCCCTTATCTTTTTC | This study |
| TCS15-C | GGGAGTTTATAGCTATGCATTAATCTCTACCTCCTGAAAAAAC | This study |
| TCS15-D | GGAGTCGACTTTCATATTGATAAGCGCTCC | This study |
| TCS16-A | GGAGGATCCATTGATGAGTGGTGTGCC | This study |
| TCS16-B | CATGACTTACACCCTAATTCATC | This study |
| TCS16-C | TAGGGTGTAAGTCATGTTTTAGAGTTTGAAATTAATATAATTCAGTATAA | This study |
| TCS16-D | GGAGTCGACATGGTCATACGAAAGCATATC | This study |
| TCS17-A | GGAGGATCCAAATGATGGACCCATGCC | This study |
| TCS17-B | CATCTATAATCTTCTTCCTTCAATTG | This study |
| TCS17-C | GGAAGAAGATTATAGATGGAATAAAAACTTTCAATATTGTAAGTATACTA | This study |
| TCS17-D | GGAGTCGACAAGTGCACCTGTAGGTTC | This study |
| **confirmation of target deletion** | | |
| TCS3-E | GTGTGTTATCAGAAACAATTGATC | This study |
| TCS3-F | GGTGCGTTAATCTTTGGTCC | This study |
| TCS4-E | GTCGAGTTAAAAGAAACATATGATAC | This study |
| TCS4-F | GGCCACACCAAATACCATAC | This study |
| TCS5-E | CATACCTGGGAGTCGTTATG | This study |
| TCS5-F | CTCTTGCAGCAGATGTTCG | This study |

| | | |
|---|---|---|
| TCS6-E | GTTTGTTTAGCTTAAGCAACCC | This study |
| TCS6-F | CAATTTGATGATGGTGTTGGTG | This study |
| TCS7-E | TTATGATACTAAGTTACTTGAAAATCG | This study |
| TCS7-F | TATCAACCCCTATAAGCCTAAC | This study |
| TCS8-E | GTGAGAATCATTGTCAATTAGAAAC | This study |
| TCS8-F | TGATCTGAAACAATTCCTGCTG | This study |
| TCS9-E | CCAACTCAAGTGATAACAAGTG | This study |
| TCS9-F | CCATCATACTCATATCATCACC | This study |
| TCS10-E | CATTAATTATGAAACAGGTCATGC | This study |
| TCS10-F | CCATTAAGTGATGCAAATCCTAC | This study |
| TCS11-E | TGCCTTAACATTTGCTTTGTATATC | This study |
| TCS11-F | CGAAAATTGGTTGGTTATCTGG | This study |
| TCS12-E | ATGACACACAAATATATATCAACGC | This study |
| TCS12-F | CTGTAATTAGTCATTTCCTATTCG | This study |
| TCS13-E | TGAGGAGAGTGGTGTAAAATTG | This study |
| TCS13-F | TGTAGTCATTTATACGAAGGGAG | This study |
| TCS14-E | ACCTGATGCACTAGATGTAAAC | This study |
| TCS14-F | GATCTAAGGTTATGTAATTGGCC | This study |
| TCS15-E | CGTTATCGTTCAATAGCACAATG | This study |
| TCS15-F | AGTGAAAGGGACAAACCAATG | This study |
| TCS16-E | ATTTATGTTAAAACCAGATGCGTC | This study |
| TCS16-F | ATTTTAAACAAGACACTACAGTCAC | This study |
| TCS17-E | GGTCATTTCTTTGGCATGCG | This study |
| TCS17-F | TAACATCATCAATGCCTTTGACG | This study |
| **in trans complementation (pHY-12, 13, 17 construction)** | | |
| TCS12-CF (S) | ATTCCCGGGAAAGAACAACTTAGCAAGTT | This study |
| TCS12-CR | GTAGGATCCTTTTCTTAATTCGATACGAA | This study |
| TCS13CPrF(HindIII) | TACAAGCTTCAGTTAAGTATTTATTTCCT | This study |
| TCS13CPrR | TCTACTTCCATAGACACTGTTATTATAC | This study |
| TCS17CF | GTGGAATTCAAATATGAATCAAAAGCAGT | This study |
| TCS17CR | TCAGGATCCTGTTAGTTCTATATTAAAAC | This study |
| **chromosomal complementation (pMADtet-13, 17 construction)** | | |
| TCS13-G | TATGGATCCAAGTGTATTTTTGTTTACCT | This study |
| TCS13-H | TAGGTCGACTAAGGTAAAACCTGTTGATA | This study |
| TCS17-G | ATGGGATCCTTGTCCCTTTTAAATATGAA | This study |
| TCS17-H | AATGTCGACGACGTTTTTGAATCAAGTG | This study |
| **SCC*mec* amplification** | | |
| mecAF | GTAGTTGTCGGGTTTGGT | 13 |
| mecAR | GGTATCATCTTGTACCCA | 13 |
| 3.0-R | CTCAGACAGCAATTTCCCG | 13 |
| ccrA-F | ACGTCAAAGTACGATGAAACAAC | 13 |
| ccrA-R | CTGACTTGTTCTCCAATGTTATCTG | 13 |
| Xsau325 | GGATCAAACGGCCTGCACA | 13 |
| attL-F | ACTTATGATACGCCTCTGCTT | This study |
| attR-R | AGAAGCTTATCATAAGTAATGAGG | This study |
| ccrA.F | TGAATGCTTCACGCTTTGTC | This study |
| ccrA.R | TTGCGTTTGTTCTCTGAACG | This study |
| **attB* mutant construction** | | |
| attB-A | CGCGGATCCCCCAGCAGCGATGTTTGTAT | This study |
| attB-B | TTTAGTTTTACTT**G**TG**G**TA**A**GC**T**TCTCCACGCATAATCTTA | This study |
| attB-C | GCGTGGAGA**A**GC**T**TA**C**CA**C**AAGTAAAACTAAAAAATTCTGT | This study |
| attB-D | AATGTCGACCATTAAGCAGTTCATAATTGTCATCA | This study |
| **attB* mutant checck** | | |
| attB-E | AAAATATTGGTATAATAAGAGG | This study |
| attB-F | AATGTCGACGCAAAATCAATTCCGAAGT | This study |
| **ccrAB mutant construction** | | |
| ccrAB-A | ATTCGCGGATCCGCAACACAGGCAATCGTATG | This study |
| ccrAB-B | GATAGCCTGTTTCTGTGCTGCAAGAGA | This study |
| ccrAB-C | CAGCACAGAAACAGGCTATCTAAGGGATTTGCAGATTCGAAGA | This study |
| ccrAB-D | TTACGCGTCGACATGCAGGTTGTTCTTGTTCATG | This study |
| **ccrAB mutant check** | | |
| ccrAB-E | AAAAGTTGGCACAAGGCATC | This study |
| ccrAB-F | GTTGGAGCTCAGGTCGATTC | This study |

Underlined, restriction site included in the oligonucleotide.

Bold, nucleotidic changes to cause silent mutations in Orfx.

Supplementary Table 3

**Supplementary Table 3. MIC (μg $mL^{-1}$) of cell wall-targeting antibiotics in CS2 medium or in TSB.**

| | **Vancomycin** | | **Bacitracin** | | **Nisin** | |
|---|---|---|---|---|---|---|
| | CS2 | TSB | CS2 | TSB | CS2 | TSB |
| Nef | 1 | 2 | 64 | > 256 | 64 | 128 |
| Nef- Δ12 | 0.25 | 0.5 | - | - | - | - |
| Nef-Δ12 (pHY-12) | 1 | 1 | - | - | - | - |
| Nef-Δ12 (pHY) | 0.25 | 0.5 | - | - | - | - |
| Nef-Δ17 | - | - | 4 | 8 | 8 | 16 |
| Nef-Δ17 (pHY-17) | - | - | 32 | 256 | 32 | 64 |
| Nef-Δ17 (pHY) | - | - | 4 | 8 | 8 | 16 |

### Supplementary Table 4

**Supplementary Table 4. Intra and interspecies transformation of distinct SCC*mec* elements in the biofilm growth conditions.**

| Donor | SCC type | Recipient | | | | | | | |
|---|---|---|---|---|---|---|---|---|---|
| | | Nef | Nef∆ comE | NefattB* | 1s | 9s | 9s∆comE | 11s | 35s |
| *S. aureus*<br>COL | I | **$3.3\times10^{-8}$** (n=1)<br>ND (n=1) | ND (n=2) | | **$1.4\times10^{-8}$**<br>± $1.1\times10^{-8}$ (n=2) | **$2.4\times10^{-8}$**<br>± $1.1\times10^{-8}$ (n=2) | | **$9.1\times10^{-8}$**<br>± $1.0\times10^{-7}$ (n=2) | **$1.7\times10^{-8}$**<br>± $1.9\times10^{-8}$ (n=2) |
| *S. aureus*<br>N315 | II | **$7.2\times10^{-8}$**<br>± $6.8\times10^{-8}$ (n=5) | ND (n=2) | ND (n=3) | **$4.6\times10^{-8}$**<br>± $4.6\times10^{-8}$ (n=4) | **$1.1\times10^{-8}$**<br>± $3.9\times10^{-9}$ (n=5),<br>ND (n=2) | ND (n=4) | **$2.3\times10^{-8}$**<br>± $1\times10^{-8}$ (n=4) | **$5.3\times10^{-8}$**<br>± $4.3\times10^{-8}$ (n=6) |
| *S. aureus*<br>N315-ΔccrAB | II<br>(ΔccrAB) | ND (n=4) | ND (n=3) | | | ND (n=3) | ND (n=3) | | |
| *S. aureus*<br>Nef | (none) | | | | ND (n=1) | ND (n=1) | ND (n=1) | ND (n=1) | ND (n=1) |
| *S. aureus*<br>35s-CoNS17 | IVa | **$9.3\times10^{-9}$** (n=1),<br>ND (n=1) | ND (n=2) | | **$5.6\times10^{-8}$**<br>± $3.1\times10^{-8}$ (n=2) | **$1.4\times10^{-8}$**<br>± $2.5\times10^{-9}$ (n=2) | | **$1.7\times10^{-8}$** (n=1),<br>ND (n=1) | **$1.1\times10^{-8}$**<br>± $8.4\times10^{-9}$ (n=2) |
| *S. aureus*<br>MW2 | IVa | **$2.1\times10^{-8}$**<br>± $1.0\times10^{-8}$ (n=2) | ND (n=2) | | **$3.4\times10^{-8}$**<br>± $4.1\times10^{-8}$ (n=2) | **$5.2\times10^{-8}$**<br>± $2.2\times10^{-8}$ (n=2) | | **$3.3\times10^{-8}$**<br>± $2.3\times10^{-8}$ (n=2) | **$3.7\times10^{-8}$**<br>± $4.5\times10^{-8}$ (n=2) |
| CoNS16 | I | **$9.6\times10^{-9}$** (n=1),<br>ND (n=1) | ND (n=2) | | **$2.1\times10^{-8}$**<br>± $1.5\times10^{-8}$ (n=2) | **$5.0\times10^{-8}$** (n=1)<br>ND (n=1) | | **$2.5\times10^{-8}$**<br>± $2.4\times10^{-8}$ (n=2) | **$5.4\times10^{-8}$**<br>± $7.3\times10^{-8}$ (n=3) |
| CoNS9 | III | **$6.2\times10^{-8}$**<br>± $6.9\times10^{-8}$ (n=2) | ND (n=2) | | **$9.7\times10^{-8}$** (n=1) | **$3.5\times10^{-8}$**<br>± $1.4\times10^{-8}$ (n=3) | | **$5.4\times10^{-8}$** (n=1) | **$3.3\times10^{-8}$**<br>± $3.5\times10^{-8}$ (n=2) |
| CoNS10 | IVa | **$1.9\times10^{-7}$**<br>± $2.6\times10^{-7}$ (n=2) | ND (n=3) | | **$5.5\times10^{-8}$** (n=1) | **$2.7\times10^{-8}$** (n=1) | | **$3.4\times10^{-8}$**<br>± $1.9\times10^{-8}$ (n=3) | **$2.9\times10^{-8}$**<br>± $2.7\times10^{-8}$ (n=2) |
| CoNS11 | IVa | **$1.6\times10^{-7}$**<br>± $1.9\times10^{-7}$ (n=2) | ND (n=2) | | **$1.5\times10^{-7}$** (n=1) | **$9.1\times10^{-9}$** (n=1) | | **$5.2\times10^{-8}$** (n=1) | **$2.5\times10^{-8}$**<br>± $2.3\times10^{-8}$ (n=2) |
| CoNS15 | IVa | **$1.2\times10^{-8}$**<br>± $1.2\times10^{-8}$ (n=2) | ND (n=2) | | **$5.25\times10^{-8}$**<br>± $2.7\times10^{-8}$ (n=2) | **$1.9\times10^{-8}$**<br>± $1.4\times10^{-8}$ (n=2) | | **$1.8\times10^{-8}$**<br>± $2.9\times10^{-9}$ (n=2) | **$3.4\times10^{-8}$**<br>± $4.4\times10^{-8}$ (n=2) |
| CoNS17 | IVa | **$2.1\times10^{-8}$**<br>± $6.0\times10^{-10}$ (n=2) | ND (n=2) | | **$4.1\times10^{-8}$**<br>± $3.3\times10^{-8}$ (n=5) | **$2.6\times10^{-8}$**<br>± $1.2\times10^{-8}$ (n=4) | | **$2.6\times10^{-8}$** (n=1) | **$2.2\times10^{-8}$**<br>± $1.7\times10^{-8}$ (n=2) |

| Donor ($Tet^R$) | | Recipient | | | | | |
|---|---|---|---|---|---|---|---|
| | | Nef | Nef∆ comE | 1s | 9s | 11s | 35s |
| Nef∆CLS2-tetR | | **$8.3\times10^{-8}$**<br>± $1.4\times10^{-8}$ (n=2) | ND (n=1) | **$2.7\times10^{-9}$**<br>± $2.9\times10^{-9}$ (n=2),<br>ND (n=2) | **$1.3\times10^{-8}$**<br>± $1.5\times10^{-8}$ (n=2),<br>ND (n=2) | **$1.0\times10^{-8}$**<br>± $1.2\times10^{-8}$ (n=3),<br>ND (n=1) | ND (n=1) |
| Nef-pT181 | | ND (n=2) | ND (n=2) | ND (n=2) | ND (n=2) | ND (n=2) | ND (n=2) |

The mean is shown with ± s.d.
n: number of independent experiments.
ND: none detected (c.a. < $10^{-9}$)

Supplementary Table 5

**Supplementary Table 5. MIC (μg $mL^{-1}$) of cefmetazole and cefoxitin in clinical isolates and lab strains in TSB.**

| | strain | Cefmetazole | Cefoxitin |
|---|---|---|---|
| MSSA recipients | Nef | 4 | 4 |
| | 1s | 4 | 4 |
| | 9s | 4 | 4 |
| | 11s | 4 | 4 |
| | 35s | 4 | 4 |
| MRSA donors | N315 | 8 | 8 |
| | MW2 | 16 | 16 |
| MR-CoNS donors | CoNS10 | 16 | 16 |
| | CoNS11 | 64 | 64 |
| | CoNS15 | 64 | 64 |
| | CoNS17 | 32 | 32 |
| Transformants (Recipient [Donor]) | 35s[N315] | 64 | 64 |
| | 35s[CoNS10] | 64 | 64 |
| | 35s[CoNS11] | 64 | 64 |
| | 35s[CoNS15] | 64 | 64 |
| | 35s[CoNS17] | 64 | 64 |
| | 1s[CoNS11] | 8 | 8 |
| | 1s[CoNS15] | 8 | 8 |
| | 9s[CoNS11] | 8 | 8 |
| | 9s[CoNS15] | 8 | 8 |
| | 11s[CoNS11] | 8 | 8 |
| | 11s[CoNS15] | 8 | 8 |

Supplementary Fig 1

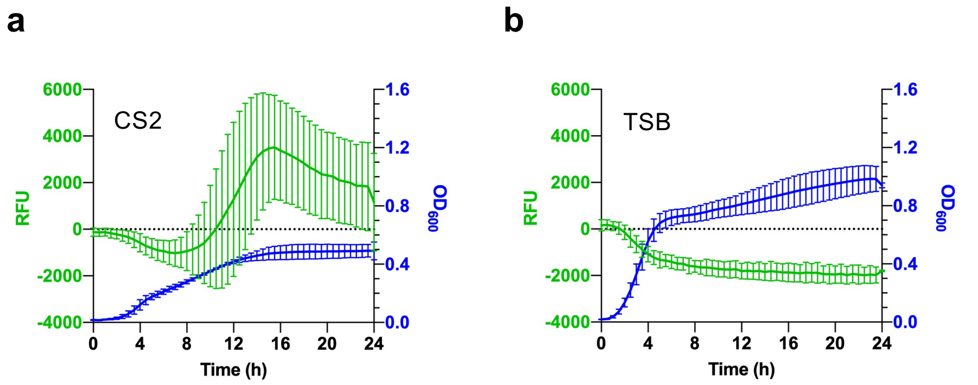


**Supplementary Figure 1. Reporter assay for *comG* promoter activity.**
Nef carrying the P*comG-gfp* reporter was grown in either CS2 medium (**a**) or TSB (**b**) with shaking. Fluorescence intensity (green line) and $OD_{600}$ (blue line) were measured every 30 min. The mean of n = 3 independent experiments is shown. Error bars represent s.d.

Supplementary Fig 2

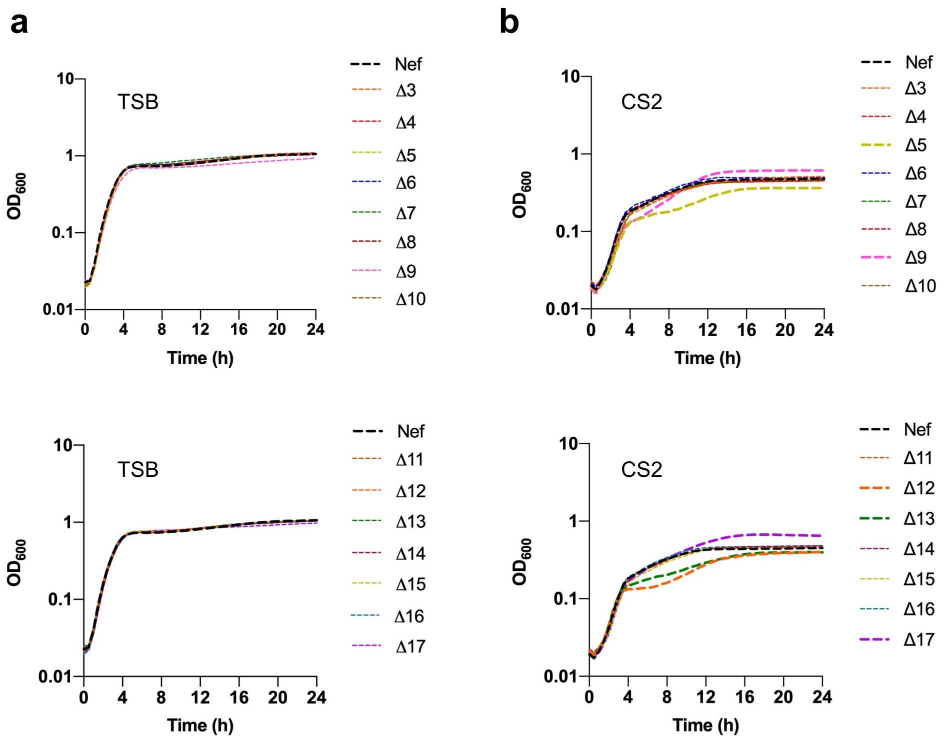


**Supplementary Figure 2. Growth curves of Nef and its derivative ΔTCS.**
Cells were grown in either TSB (**a**) or CS2 medium (**b**) with shaking. $OD_{600}$ was measured every 30 min. The mean of n = 3 independent experiments is shown. Error bars are omitted for clarity.

Supplementary Fig 3

a

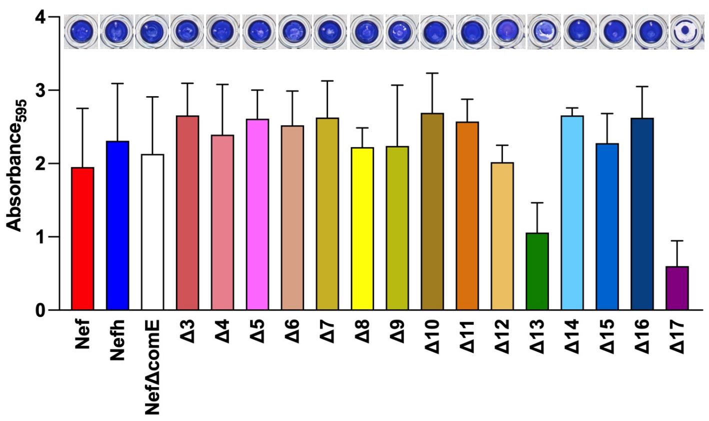


**Supplementary Figure 3. Biofilm formation is impaired in the TCS13 and TCS17 mutants.**

(**a**) Nef and its derivatives were statically grown in CS2 medium in a 96-well plate for 24 h.

(**b**) Time-course of biofilm development in Nef and its derivatives. The cells were statically grown in CS2 medium in a 6-well plate for up to 4 days. (**b, middle)** The biofilms were stained with crystal violet and the absorbance at 595 nm was measured. The mean of n = 2-3 independent experiments is shown. Error bars represent SD. (**b, right**) CFU of the cells after 24 h in the biofilm.

b

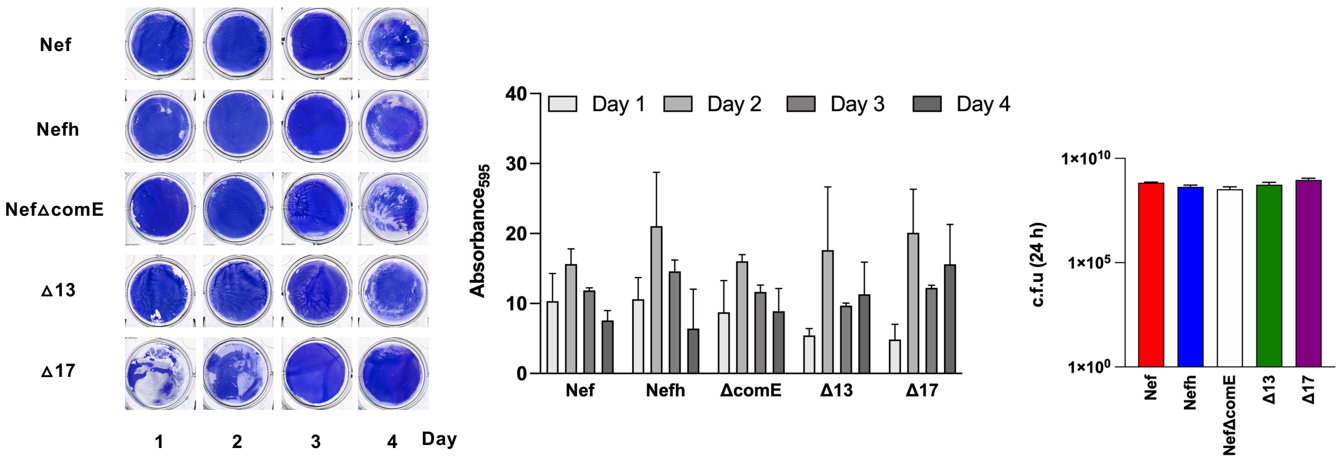

Supplementary Fig 4

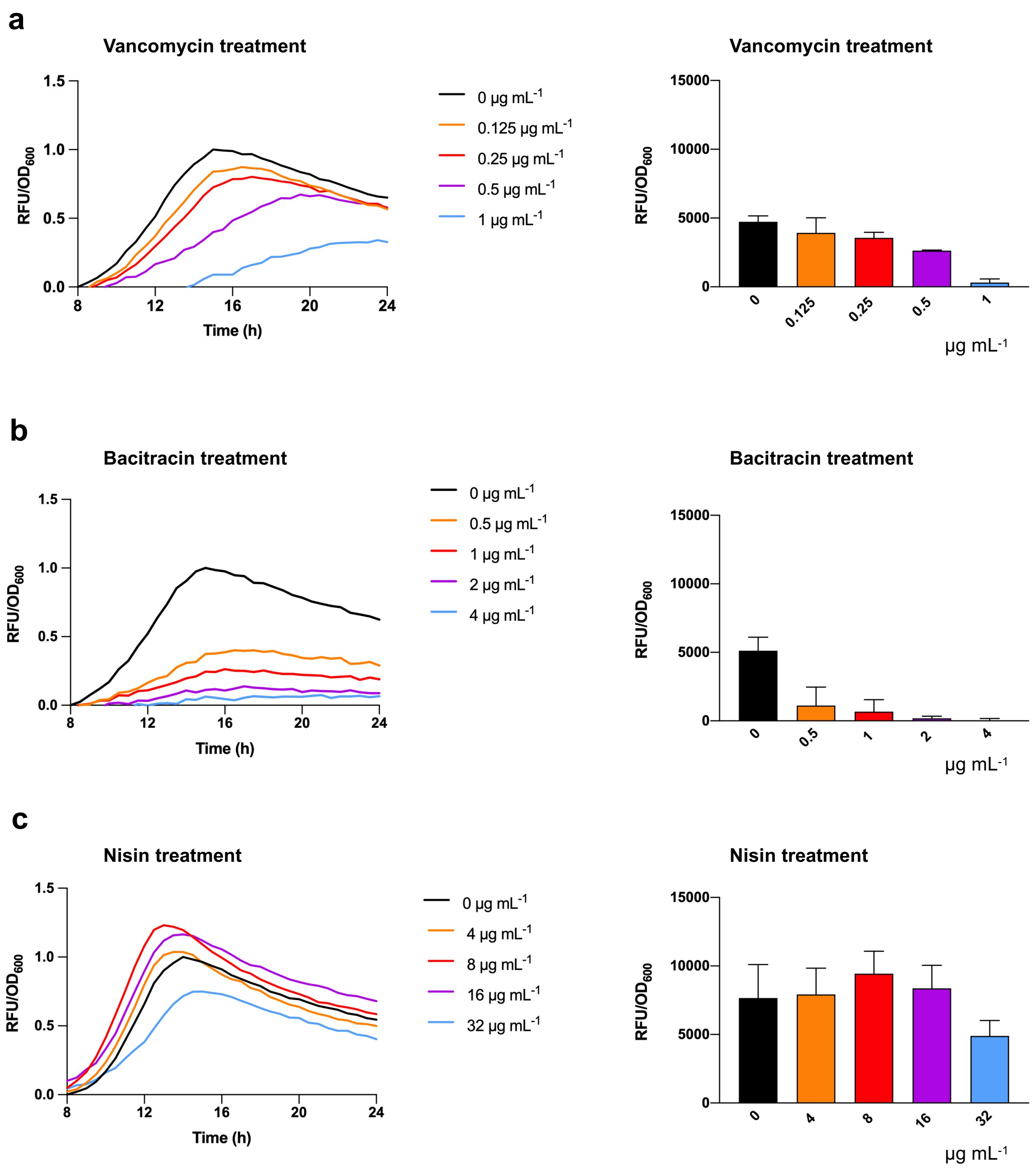


**Supplementary Figure 4. *comG* promoter activity is affected by cell wall-targeting antibiotics.**
Nef-GFP was treated with subinhibitory concentrations of vancomycin (**a**), bacitracin (**b**), or nisin (**c**). Cells were grown in CS2 medium with shaking for 24 h. Fluorescence (RFU) and $OD_{600}$ were measured every 30 min. Data shown are either relative RFU/$OD_{600}$ values (**left panels**) or increases in RFU/$OD_{600}$ values during 8-24 h of growth (**right panels**).

Supplementary Fig 5

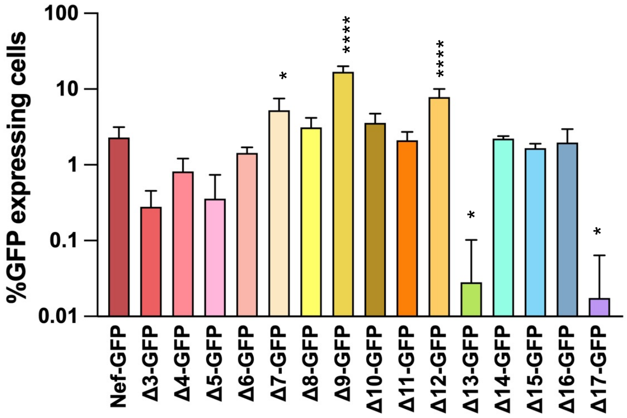


**Supplementary Figure 5. *comG* promoter activity is affected by multiple TCSs in biofilm.** Nef and its derivative ΔTCSs carrying the P*comG-gfp* reporter were statically grown in CS2 medium in biofilm growth conditions for 3 days. The percentage of GFP-expressing cells was determined by fluorescence microscopy. The mean of n = 3-7 independent experiments is shown. Error bars represent s.d. Statistical significance was determined by one-way ANOVA with Tukey's multiple comparison test. *P<0.05, ****P<0.0001.

Supplementary Fig 6

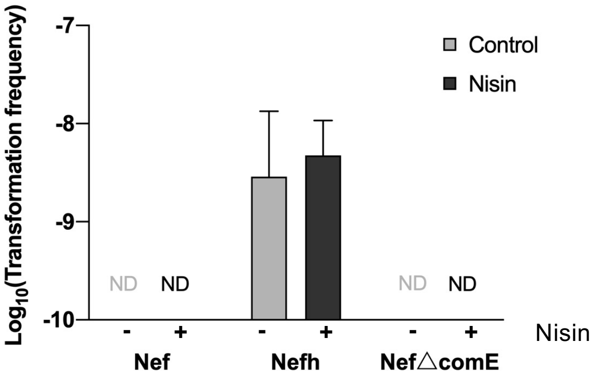


**Supplementary Figure 6. Nisin does not induce natural transformation in Nef**.

Nef and its derivatives were grown in CS2 medium with or without nisin (8 μg $mL^{-1}$). Transformation frequencies were determined after 10 h of planktonic growth. The transformants were selected by tetracycline. The mean of two independent experiments is shown with s.d. ND < $10^{-9}$.

Supplementary Fig 7

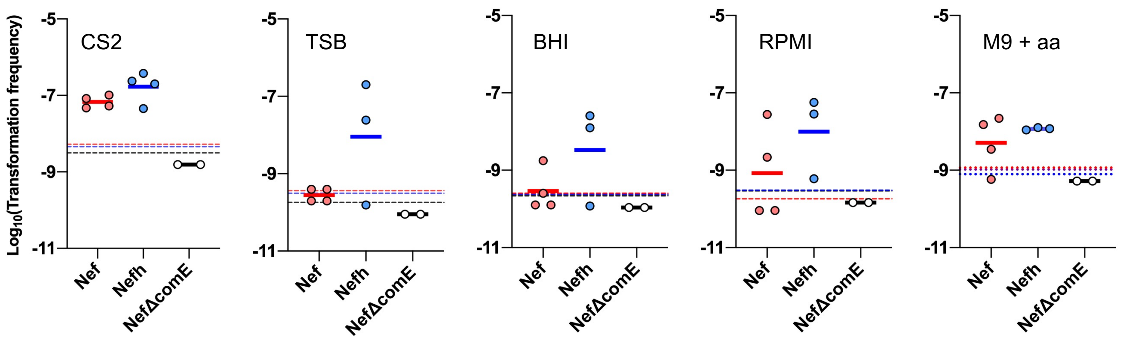


**Supplementary Figure 7. Nutrient-poor culture media are preferable for natural transformation under biofilm growth conditions**. Nef, Nefh, and NefΔcomE were statically grown in different growth media, including CS2, TSB, BHI, RPMI, and M9 supplemented with amino acids (M9 + aa). Transformation frequencies were determined at day 3. Dotted lines represent the detection limit of the strains in each growth medium. Data points represent independent experiments. The mean is shown.

Supplementary Fig 8

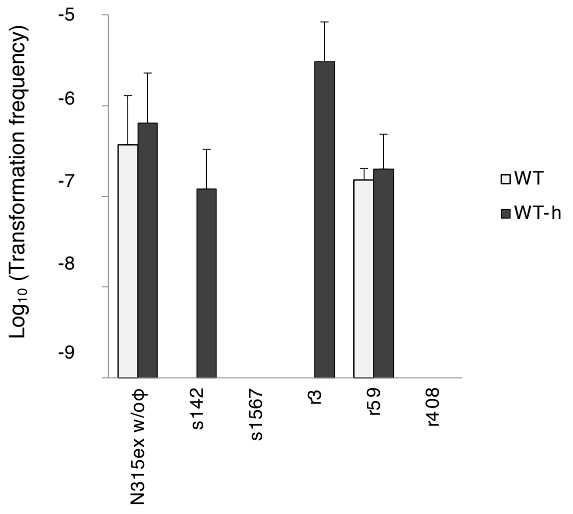


**Supplementary Figure 8. Transformation frequencies of clinical isolates.** Wild-type (WT) *S. aureus* and its derivative carrying the SigH expression plasmid pRIT-sigH (WT-h) were tested for transformation after 2 days of static growth in CS2 medium. The transformants were selected by tetracycline. The mean of n = 2- 3 independent experiments is shown. Error bars represent s.d.
